# Cathodoluminescent and Characteristic X-ray-emissive Rare-Earth-doped Core/Shell Immunolabels for Spectromicroscopic Analysis of Cell Surface Receptors

**DOI:** 10.1101/2024.03.20.585848

**Authors:** Sebastian Habermann, Lukas R. H. Gerken, Mathieu Kociak, Christian Monachon, Vera M. Kissling, Alexander Gogos, Inge K. Herrmann

## Abstract

Understanding the localization and the interactions of biomolecules at the nanoscale and in the cellular context remains challenging. Electron microscopy (EM) as a non-Abbe limited technique gives access to the cellular ultra-structure yet results in grey-scale images and averts unambiguous (co-)localization of biomolecules. Multimodal nanoparticle-based immunolabels for correlative cathodoluminescence electron microscopy (CCLEM) and energy-dispersive X-ray spectromicroscopy (EDX-SM) are presented. The single-particle STEM-cathodoluminescence (CL) and characteristic X-ray emissivity of sub-20 nm lanthanide-doped nanoparticles were exploited as unique spectral fingerprints for precise localization and label identification. To maximize the nanoparticle brightness, lanthanides were incorporated in a low-phonon host lattice and separated from the environment using a passivating shell. The core/shell nanoparticles were then functionalized with either folic (terbium-doped) or caffeic acid (europium-doped). Their potential for immunolabeling was successfully demonstrated using HeLa cells expressing different surface receptors that bind to folic or caffeic acid, respectively. Both particle populations showed single-particle CL emission along with a distinctive energy-dispersive X-ray signal, with the latter enabling colour-based localization of receptors within swift imaging times well below 2 mins per µm^2^ while offering high resolution with a pixel size of 2.78 nm. Taken together, these results open a route to color immunolabelling based on electron spectromicroscopy.

**Table of Contents:** 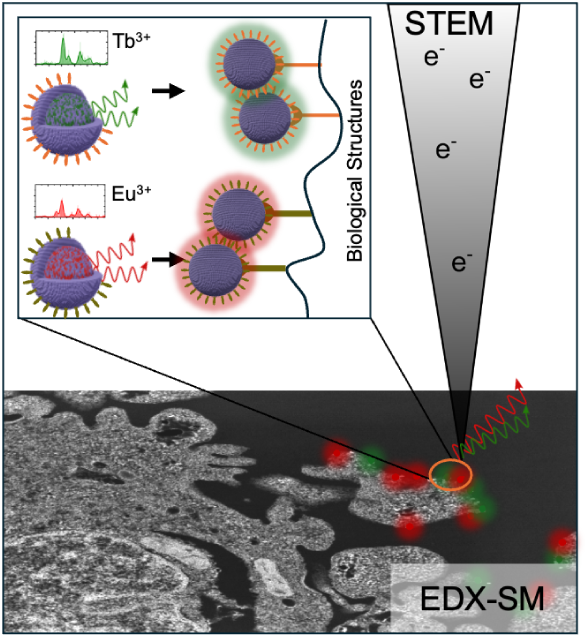
Small (sub-20 nm) lanthanide-doped nanoparticles were successfully utilized in electron microscopy to label biological structures and contextualize them in the cell’s ultrastructure. Leveraging unique energy-dispersive X-ray signatures, the nanoparticles’ location and doping-identity was easily and fast retrieved, demonstrating the methods’ potential to (co)-localize labels while supplying a holistic impression of the underlying processes, as entire cells could be mapped.

## 1 Introduction

The microscopic exploration of biological and material structures has been a cornerstone in advancing scientific knowledge. By revealing the molecular organization of e.g. lipid membranes, cytoplasmic components or specific biomolecules, ultrastructural analysis provides an ultimate platform to investigate structure-function relationships. [1] Over the decades, various imaging techniques have emerged, providing deeper insights into the intricate properties of these structures. Although light and fluorescence microscopy are commonly used in biological research to gain functional insights, their achievable resolution is limited by the wave-like behaviour and diffraction events of visible light to the so-called Abbe limit. [2] Emerging super-resolution microscopy (SRM) techniques such as stimulated emission depletion (STED) or stochastic optical reconstruction microscopy (STORM) offer higher resolutions by specifically exploiting the optical properties of labels. [3,4] However, the reliance on fluorescent labels results in a lack of ultrastructural context, as most cell components remain unlabeled and therefore invisible. [5] Electron microscopy (EM) remains the method of choice for ultrastructural analysis, enabling the visualization of various cellular components ranging from nanometer-sized ribosomes to cell-spanning structures such as microtubules, mitochondria, vesicles or the nucleus. [6–8] However, EM based on electron density yields low-contrast grayscale images where pinpointing the distributions of specific proteins is often elusive. While correlative light and electron microscopy (CLEM) approaches have been developed, they necessitate intricate procedures and meticulous alignment of datasets. [5]

Recently, correlative cathodoluminescence electron microscopy (CCLEM) and electron-dispersive X-ray spectromicroscopy (EDX-SM) denoted as ColorEM have gained considerable attention due to their high spatial resolution and the ability to directly probe optical and electronic properties at the nanoscale within the electron microscope. [9] Yet, a significant challenge in the field has been to develop suitable luminescent labels that can effectively and specifically enhance contrast in CCLEM or EDX-SM and enable the unambiguous identification and localization of specific proteins in biological specimens within reasonable imaging times. [10]

With their unique electronic configurations and sharp ff-transition-luminescence lines, rare-earth elements offer compelling prospects as luminescent materials. When doped into host lattices, these elements can produce luminescent centers that exhibit exceptional stability, unique emission patterns with narrow line widths, and long lifetimes. [11,12] Recently, the development of rare-earth element-doped nanocrystals has opened up promising avenues in the realm of CCLEM as well as EDX-SM. [12,13] When tailored as immunolabels, these nanocrystals can potentially revolutionize the visualization capabilities in CCLEM and EX-SM by enabling swift multi-color imaging. However, reliable single-particle cathodoluminescence (CL) emission from sub-20 nm nanocrystals has shown to be challenging due to the limited signal generation volume, low stability, and quenching events within and on the crystal’s surface. A first feasibility study was reported by Prigozhin *et al.* in which the CL response of silicon wafer-deposited sub-20 nm rare-earth-doped nanocrystals was investigated using a scanning electron microscopy (SEM) set-up. [10,14] It was shown that secondary electron images and simultaneously acquired CL signals can be correlated, as sufficient signal-to-noise ratios were achieved, enabling pinpointing the crystal’s location based on its CL emission. Thereby, sodium gadolinium fluoride (NaGdF_4_), a well-known host material, has been exploited, since it exhibits favourable properties, such as a low phonon energy, which can reduce non-radiative relaxations and thereby enhance luminescence efficiency. [15] These nanocrystals not only harness the advantages of the host lattice but also encapsulate the distinctive luminescent properties of rare-earth dopants. However, the translation of these findings to biological samples has yet to be demonstrated, as the particles were neither phase-transferred (functionalized) nor introduced into biological sample conditions (e.g. embedding and staining). [16]

CL-enabled immunolabelling of cells was investigated by Keevend *et al.* by comparing and bench-marking the membrane-specific receptor interactions of folic acid-decorated CL-active YPO_4_:Tb nanoparticles with their immunogold counterparts. Labelling and blocking experiments proved the specificity and feasibility of the nanoparticles as CL labels, and the EM image was successfully correlated with the cathodoluminescence signal. However, the size of the investigated particles (50-70 nm) limited their use in the field of immunolabeling. [17] Moreover, these investigations have been limited to single dopant populations and overall CL intensity, not taking advantage of the rare-earths’ multi-colour potential and their unique ff-transition emission patterns. In 2021, Swearer *et al.* demonstrated cathodoluminescence spectra with assignable transitions from differently doped single BaYF_5_:Ln^3+^ particles, however, operating in ideal imaging conditions on a TEM grid instead of biological samples, and requiring unfeasibly long acquisition times of 30 s at the centre of each nanoparticle to achieve the spectral information required for reliable dopant identification. [18] Recently, Rehman *et al.* utilized lanthanide-based cathodophores as an unspecific staining of a cell’s surface. In this SEM-based multi-color approach, spectral filters were introduced, simplifying the dual dopant recognition on the outer cell membrane, but veiling nonlocal excitations through secondary electrons were found and the obstacles associated with epoxy-embedding, required for intracellular ultrastructure assessment, were neglected. [19,20]

Alternatively to CL, characteristic X-ray emission may be leveraged with EDX-SM for label identification and characterization of the elemental distribution within samples. Work by Scotuzzi *et al.* demonstrated its feasibility using quantum dots and gold-based immunolabels. [21] Their contrasting elemental compositions allowed for the labels’ co-localization, yet differences in morphological parameters of these labels (especially size and shape) are known to result in different penetration properties, which needed to be addressed during immunolabelling protocols. [22,23] While especially appealing for the fluorescent label-free assessment of elemental distributions of exogenous (e.g. nanoparticle label) and endogenous constituents within biological samples, the acquisition times of several hours severely limit the impact of EDX-SM ColorEM and constrains its use to a niche. The elemental mapping of endogenous components is occasionally applied, such as for the investigation of iron-deficient mitochondria by Hoes *et al.* or used by Giepmans *et al.* to determine the granule content in human type 1 diabetes cells. [24,25] However, despite the capabilities of EDX-SM to probe even light (low-Z) elements, biological samples are not routinely mapped, and the potential of rare-earth-based immunolabels as substitute for the gold standard remains untouched. [9] Hence, new approaches are needed to increase the nanoparticle label brightness and characteristic X-ray emission yield, while meeting the morphological constraints imposed by the immunolabelling application as well as allowing reasonable imaging conditions.

In this work, we demonstrate the effectiveness of using cathodoluminescent and energy-dispersive X-ray spectroscopyemissive immunolabels for ColorEM-based nanoscopy. Our approach comprises the synthesis of sodium gadolinium fluoride nanoparticles with a core/shell structure, being less than 20 nm in size, and doped with the rare-earths europium or terbium. To meet the dual requirements of the small size necessary for precise immunolabelling while guaranteeing sufficient signal generation (constrained by particle size), we exploited a strategy involving a protective shell to amplify the CL signal. [19,26–29] By substituting the lanthanides within a host matrix, an adjustable platform is provided, which creates labels that are indistinguishable from their outer surface while the difference between the particle populations is reduced to the core’s doping. We further introduced high concentrations of rare-earth elements, leveraging the principle that the probability of CL and EDX-SM signals is proportional to the number of atoms in the particle. [19,27] To demonstrate immunolabelling capabilities, we functionalized the nanocrystals doped with terbium and europium with folic acid and caffeic acid, respectively. These compounds, intended for surface receptor targeting, allowed us to assign different targeting moieties to each rare-earth label, paving the way to ColorEM nanoscopy. [17,30,31]

## 2 Morphological and Compositional Characterization

Core/shell sodium gadolinium fluoride (NaGdF_4_) nanoparticles were prepared using a thermal decomposition process (Figure 1A). Initially synthesized nanoparticle nuclei were enlarged using a hot-injection protocol, and the grown particle core was subsequently wrapped with an undoped shell, yielding a core/shell nanoparticle architecture. Europium and terbium, both neighbouring gadolinium in the periodic table, were used as dopants, acting as unique recognizable centres in the host matrix. The nanoparticle size and shape evolution was monitored based on bright-field transmission electron microscopy (TEM) (europium: Figure 1B, a-c; terbium: Figure 1B, d-f). Both particle types exhibited diameters at their shortest axis of d_TEM_ *≈* 9 nm for the nuclei and d_TEM_ *≈* 13 nm for the doped-core stages (Figure 1C). The core/shell nanoparticles showed mean average particle diameters of d_NaGdF4:Tb_ = 19.7 *±* 4.1 nm and d_NaGdF4:Eu_ = 17.9 *±* 2.8 nm (*±* SD; Figure 1C). Thereby, the nanoparticle size met the strict size constraints dictated by the compromise between immunolabelling application and signal generation volume, which limited the particles’ maximum diameter to d_max_ *≤* 20 nm. [14,32–35] The size growth of both core/shell nanoparticle types was accompanied by a step-wise shape harmonization, resulting in particle populations with uniform hexagonal particle outlines and narrow size distributions. The similar size and shape laid the structural foundation for comparable diffusion (see Suppl. Figure S1) and penetrating properties important for ensuring equal preconditions in immunolabelling applications. [34,36,37] The EDX-SM maps of single core/shell nanoparticles (Figure 1D) showed the rare-earth dopants and gadolinium (as a proxy for the host matrix) atoms forming a homogeneous phase without noticeable crystallographic segregation. The nanoparticle cross-sections (yellow arrows in Figure 1D) revealed gadolinium being present throughout the entire diameter range, while the rare-earths showed an enrichment in the particle center (Figure 1E). Given that EDX-SM is a 2D projection of the particle, this observation is in line with the synthetic setup, in which solely the nuclei and the core particle were considered to be dopant-bearing, and strongly supports the existence of the desired core/shell architecture of the nanoparticles. The rare-earth content of the particles was determined using inductively coupled plasma optical emission spectroscopy (ICP-OES), showing that high doping concentrations close to the nominal concentration (40%) were achieved for both particle types (*≈*38.7% for europium and *≈*34.4% for terbium; Figure 1F). Together with the nearly identical core sizes, both particle types share a rather high amount of CL- and EDX-SM-response generating atoms in their cores, whose numbers are in line with recent trends in the field of lanthanide-doped nanomaterials and cathodophores. [19,27] The final core/shell nanoparticles exhibited a total rare-earth content of *≈*17.3% and *≈*14.5% for europium and terbium, respectively (Figure 1F).

**Figure 1:**
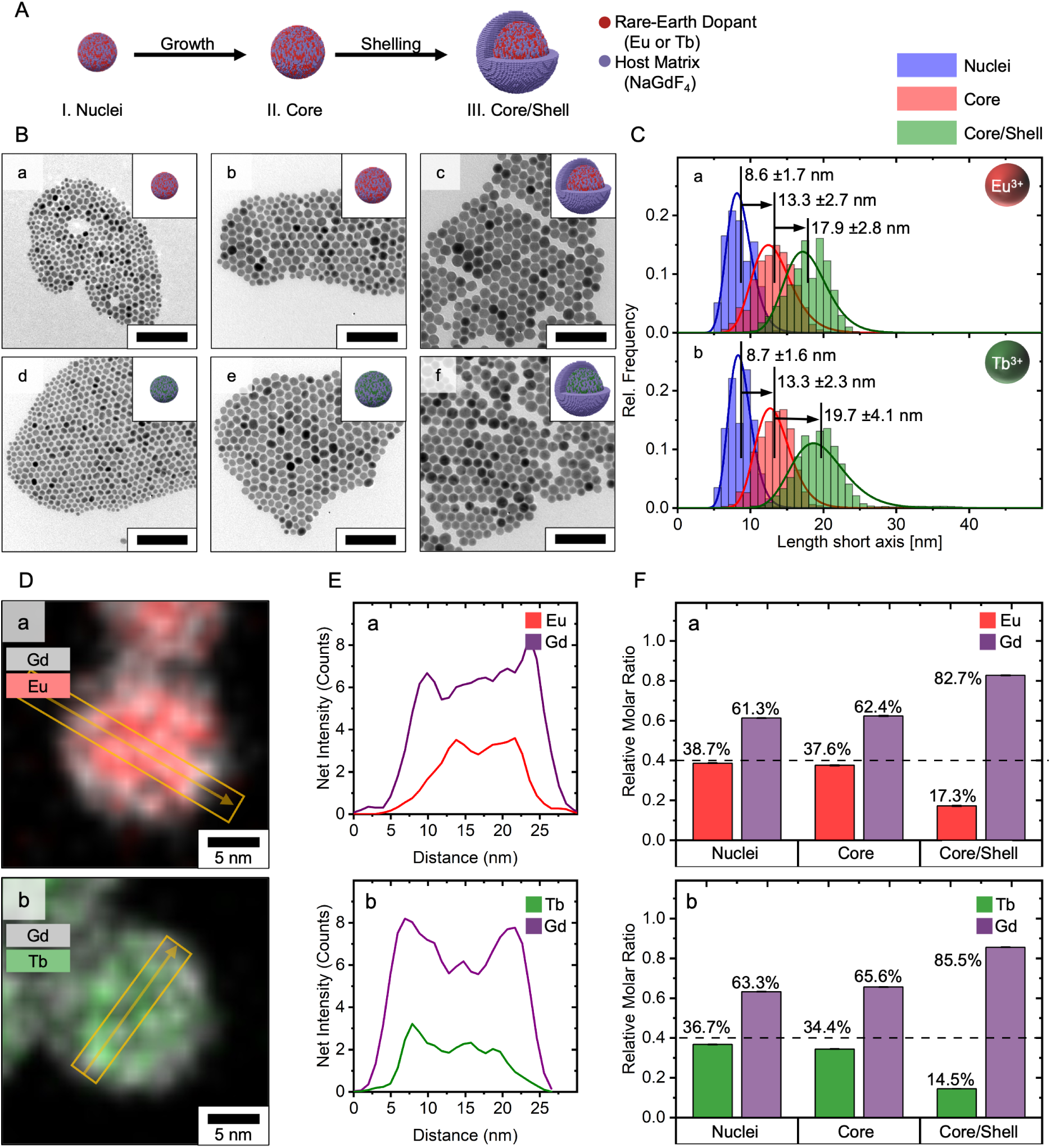
Nanoparticle Morphology and Composition. (**A**) Schematic representation of the three-step particle preparation process, starting from the growth of small particle nuclei to a subsequent coating of the particle core using an undoped shell. (**B**) Bright-field transmission electron microscopy (TEM) images showing the individual development stages of europium(**a-c**) and terbium-doped nanoparticles (**d-f**). Scale bar: 100 nm. (**C**) Corresponding particle size distribution and average particle diameter along the short axis are shown alongside *±* SD error and Lognormal fit. (**D**) Energy dispersive X-ray spectromicroscopy (EDX-SM) of differently-doped nanoparticles. (**E**) Net intensity analysis of EDX-SM cross-sections displayed in (**D**) as orange arrows. (**F**) Rare-earth content determined using inductively coupled plasma optical emission spectroscopy (ICP-OES) normalized on the total rare-earth content.

Thus, based on the physicochemical characterization, the prepared nanoparticles match the desired design requirements in size, architecture and elemental composition.

## 3 Spectral Characterization of Doped-Nanoparticles

For spectral characterization, bulk measurements of the core/shell particles were conducted, aiming to assess their response emission of low-energy CL- and high-energy X-ray-photons upon electron irradiation. The CL emission spectrum of the europium-doped core/shell nanoparticles (Figure 2C, a) was dominated by three distinctive electronic transitions originating from the rare-earth’s ^5^D_0_ (first excited) state. The prominent signals located at 591 nm and 615 nm were attributed to the ^5^D_0_ *→*^7^ F_1_ and ^5^D_0_ *→*^7^ F_2_ transitions, respectively. Furthermore, the transition at 696 nm was assigned to the ^5^D_0_ *→*^7^ F_4_ transition, and completed the characteristic emission pattern of europium as often described in the literature. [14,15,38,39]. The CL emission spectrum of the terbium-doped core/shell nanoparticles (Figure 2C, b) mainly revealed a pattern composed of four signals. All of them are known for terbium and found to originate from its first excited state (^5^D_4_) to the ^7^F*J* ground state levels: ^5^D_4_ *→*^7^ F_6_(489 nm); ^5^D_4_ *→*^7^ F_5_(542 nm); ^5^D_4_ *→*^7^ F_4_(585 nm) and ^5^D_4_ *→*^7^ F_3_(620 nm). [40,41] Both CL spectra were in strong agreement with publications investigating both the lanthanide’s photoluminescence (PL) and CL. Furthermore, the individual spectra were in accordance with PL spectra recorded (see Suppl. Figure S2), thereby confirming the host-independent emissive behaviour of the respective luminescent centre. [14] Each dopant featured unique transition patterns (locations paired with intensities) usable for their precise identification. Here, the most intense transitions (Eu: *→*^7^ F_1_,_2_; Tb: *→*^7^ F_5_,_6_ *∨* _4_) serve as important anchors, since they showed sharp non-interfering signals. Thereby, the nanoparticles’ CL response offered a suitable tool for reliable label recognition.

**Figure 2:**
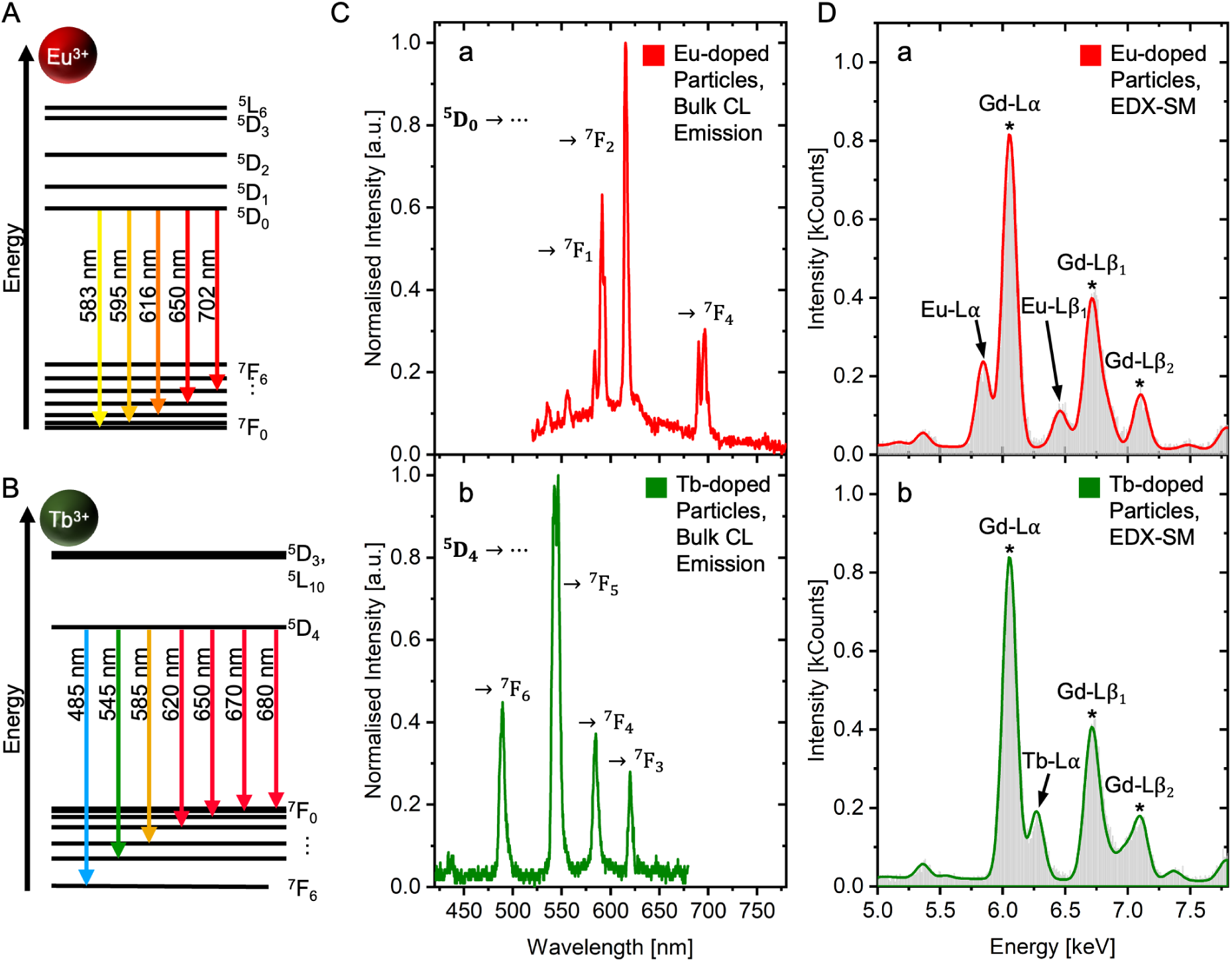
Spectral Characterization. (**A, B**) Schematic illustration of energy levels of europium and terbium ions with the indication of important electronic transitions. [38] (**C**) Cathodoluminescence (CL) response of core/shell nanoparticles (**a:** Eu^3+^-doped; **b:** Tb^3+^-doped) under irradiation with 3 kV electron beam. Important electronic transitions are labelled. a.u.: arbitrary units. (**D**) Core/shell particles’ characteristic EDX-SM spectra (**a:** Eu^3+^-doped; **b:** Tb^3+^-doped) with the rareearth’s Lα and Lβ lines highlighted.

The EDX-SM spectra of both particle types in bulk are shown in Figure 2D. Both spectra were dominated by a common signal at 6.05 keV, which was in accordance with literature assigned to the Gd-Lα lines of the rare-earth’s host material. [42] Additional gadolinium lines were visible at 6.71 keV (Gd-Lβ_1_) and 7.10 keV (Gd-Lβ_2_), building the background alongside europium and terbium. Sodium (Z=11) and fluorine (Z=9), atoms with a fundamentally different electronic structure compared to the rare-earths (Z=63-65), were not expected to show interfering signals within the sliced energy range. [42] The spectrum of the europium-doped nanoparticles showed two additional lines located at 5.84 keV (Eu-Lα) and 6.46 keV (Eu-Lβ_1_). The same applied to the terbium-doped nanoparticles, where characteristic Tb-Lα lines were detected at 6.27 keV and a broadening of the gadolinium signal at *≈*7.00 keV was attributed to the Tb-Lβ_1_ line with its theoretical value of 6.97 keV. Since the dopants and gadolinium share adjacent atomic numbers, their inner shell energy levels are similar. However, as demonstrated in Figure 2D, the EDX-SM energy resolution was sufficient to distinguish the individual elements, enabling the recognition of europium and terbium next to the host material. The respective signal intensities were the consequences of the line transition probabilities and the total number of introduced rare-earth centres in the core/shell nanoparticles. Based on the bulk measurements, both CL and EDX-SM spectra of the doped nanomaterials showed distinctive features, which allowed precise identification of the luminescent centre. Using CL, the most intense electronic transitions for Eu and Tb did not overlap and the full spectrum showed recognisable patterns, which can be used to support the identification, if a high signal quality is given. Pivotal for EDX-SM were the rare-earths’ Lα lines, which lined up every *≈*0.2 keV with increasing atomic number and were still easily distinguishable and assignable to the dopant. In the event of interference or overlapping with other elements in the sample, the Lβ lines could be included in the determination, however, as both particle populations were mono-doped, the signal was qualified for reliable deconvolution to the respective dopant. Taken together, both responses bear the potential to be exploited for ColorEM-recognition.

## 4 Surface Functionalization: Introducing Targeting Moiety

For immunolabelling application, the nanoparticle’s surface was modified, and the targeting moieties folic acid (FA) and caffeic acid (CA) were attached. Folic acid was selected as it is a natural folate receptor (FR) agonist and enabled effective targeting of FR-positive HeLa cells, when immobilized on nanoparticle surfaces. [17,43] Caffeic acid was chosen as second functionalization, since it shares a catechol-moiety that is crucially involved in receptor binding with βadrenergic receptor-agonists (e.g. epinephrine, isoetharine), and because its derivates such as caffeic acid phenethyl ester have been confirmed to bind to β-adrenergic receptors. [31,44–46] Furthermore, caffeic acid ethanolamide (CAEA), structurally closely related to caffeic acid, was shown to interact with the TGF-β_1_ receptor. [47] To facilitate the coordination of both ligands, the nanoparticles were first transferred into polar media. Afterwards, a primary amine-containing ligand was introduced to which the desired ligand was attached using an EDC/NHS coupling protocol (see Figure 3A). Thereby, a compact architecture was achieved, reducing the distance between particle and ligand, and as a result, increasing the label’s localisation precision. [26] For each nanoparticle type with its specific dopant one functionalization was assigned. Caffeic acid was arbitrarily assigned to europium-doped particles and folic acid to terbium-doped particles. Surface functionalization was confirmed using Fourier Transform Infrared spectroscopy (FTIR) and thermogravimetric analysis (TGA). The final ligand-capped stage was also examined using ultraviolet-visible (UV/Vis) spectroscopy.

**Figure 3:**
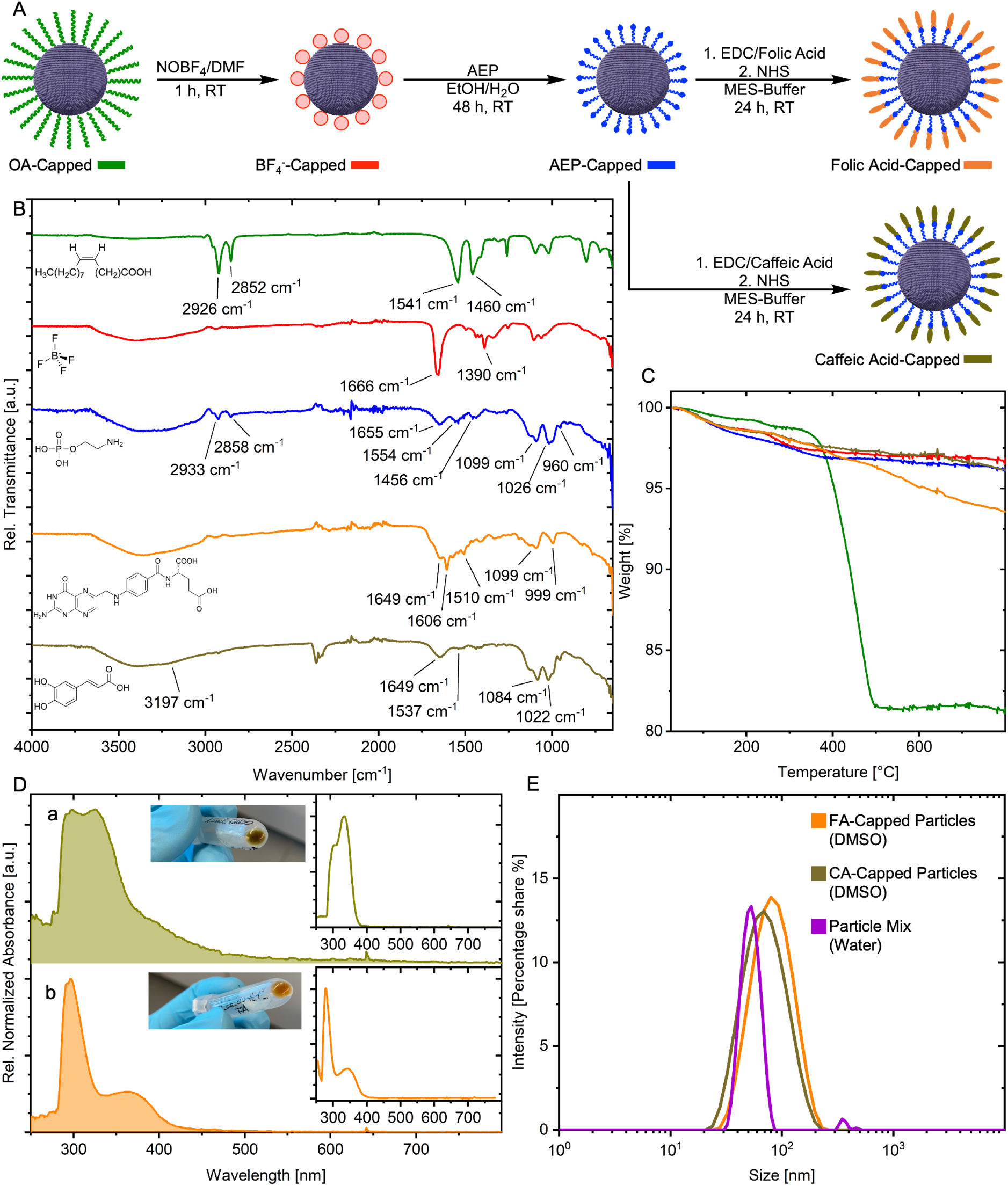
Surface Functionalization. (**A**) Schematic illustration of the surface chemical evolution and introduction of a colour code for surface functionalizations. OA Oleic Acid (green); BF4: Tetrafluoroborate (red); AEP: 2-Aminoethyl dihydrogen phosphate (blue); Folic acid (orange); Caffeic acid (orange-brown). (**B**) Fourier-transform infrared spectroscopy (FTIR) spectra recorded at each functionalization step with the modifications’ chemical structure displayed next to the corresponding signal and vibrations highlighted. a.u.: arbitrary units. (**C**) Mass loss of the nanoparticles for each functionalization step, determined using thermogravimetric analysis (TGA). (**D**) Absorption spectra of caffeic acid-capped nanoparticles (**D, a**) and folic acid-capped particles (**D, b**) were assessed through ultraviolet-visible (UV-Vis) spectroscopy. Absorption of free organic molecules shown in graph insets. a.u.: arbitrary units. (**E**) Hydrodynamic radius assessed by dynamic light scattering (DLS) of final functionalization and cell experiment-relevant mixture. FA: Folic acid, CA: Caffeic acid.

The FTIR spectrum of the as-prepared core/shell nanoparticles (Figure 3B) showed distinct signals located at 2926 cm*^−^*^1^ and 2852 cm*^−^*^1^, which indicated an alkyl component with its ν_as_(-CH_2_-) and ν_s_(-CH_2_-) vibrations modes. [48] Further, distinct signals at 1541 cm*^−^*^1^ and 1460 cm*^−^*^1^ were assigned to the ν_as_(COO^–^) and ν_s_(COO^–^) vibration modes of carboxylate groups, indicating a deprotonated terminal carboxylic acid and by this confirming the presence of oleic acid as initial surface capping. [48,49]

Subsequently, nitrosyl tetrafluoroborate (NOBF_4_) was used to substitute oleic acid and provide an easily exchangeable precursor for future modifications in polar media. After the reaction, the oleic acid-attributed vibrations located at 2800 cm*^−^*^1^ to 3000 cm*^−^*^1^(-CH_2_-) and 1400 cm*^−^*^1^ to 1600 cm*^−^*^1^(COO^–^) vanished, confirming the removal of oleic acid from the particle’s surface. Novel vibrations emerged at 1666 cm*^−^*^1^ and 1390 cm*^−^*^1^, which were found to originate from the ν(C=O) and δ*_s_*(CH_3_)N of the dimethylformamide’s (DMF) amide (see Suppl. Figure S3). [50,51] This coordination of solvent molecules on the particle surface was in agreement with literature reports for this procedure. [51] Vibrations of the tetrafluoroborate anion (BF_4_^–^) could not be unambiguously assigned due to the interfering signals of DMF around *≈*1000 cm*^−^*^1^ to 1100 cm*^−^*^1^.

After transfer into a polar medium, 2-aminoethyl dihydrogen phosphate (AEP) as terminal amine-bearing linker was introduced. The FTIR spectrum revealed the loss of previously dominant DMF vibrations and showed new vibrations located around 1300-900 cm*^−^*^1^. The complex signal was matched to ν(P—O) vibrations of the AEP’s phosphate group (PO_4_^3–^) and was in accordance with literature reports. [52,53] Apart from the particle-coordinating phosphate, and the small alkyl constituent indicated by the vibrations between 3000 - 2800 cm*^−^*^1^ and 1456 cm*^−^*^1^, the vibration at 1554 cm*^−^*^1^ was assigned to a terminal amine. [54–56]

Enabled by the terminal amine, the particles that were functionalized using FA that revealed a characteristic amide bond peak at 1649 cm*^−^*^1^. [57,58] Additionally, the signals located at 1606 cm*^−^*^1^ and 1510 cm*^−^*^1^ were assignable to vibrations related to the phenyl and pterin rings of FA. [59] Moreover, the bending vibration of the terminal amine, located at 1554 cm*^−^*^1^, disappeared after FA functionalization. In comparison to this, the CA functionalization showed a more prominent vibration at 1649 cm*^−^*^1^ and a weaker one at 1537 cm*^−^*^1^. These were assigned to the amide I and II bands, respectively. [58,60] In addition, two shoulders were detected in the broad signal between 3500 - 2800 cm*^−^*^1^, which were characteristic for O—H stretching vibrations of the catechol moiety of caffeic acid. [61] After confirming the surface modifications using FTIR, TGA was conducted to quantitatively assess chemical mass changes. As shown in Figure 3C, the as-prepared nanoparticles exhibited a mass loss of 17% in a temperature range of 280 *^◦^*C*→*540 *^◦^*C. After phase transfer, the mass loss was 1%, which was mainly attributed to surface-bound DMF and supports the chemical changes indicated by FTIR. After ligand exchange and functionalization towards AEP and CA, no significant changes were detected. The FA-capped particles, however, showed a mass loss comparable to the folic acid reference (see Suppl. Figure S4), indicating successful functionalization with FA.

Facilitated through the inherent absorption behaviour of both folic and caffeic acid, the final ligand modification was proven by ultraviolet-visible (UV-Vis) spectrophotometry (Figure 3D). As AEP exhibited no absorption throughout the measured range (data not shown), the colour-less AEP-capped nanoparticles served as a silent precursor for the absorbing ligands. After the reaction, a colour change was visible for both functionalizations accompanied by changes in their absorbance spectrum. CA functionalized nanoparticles showed increased absorbances peaking at 299 nm and 325 nm. The CA reference showed similar signals, with the most intense absorption at 334 nm and a shoulder around 305 nm. The absorptions can be referenced to the *π → π^∗^* transition from the caffeic moiety towards the carbonyl and the n *→ π^∗^* transition within the carbonyl, respectively. [60] The FA-coordinated particles showed bathochromically shifted absorptions compared to the reference spectrum with absorption peaks located at 297 nm and 366 nm. These originate from the FA-characteristic *π → π^∗^* and n *→ π^∗^* transitions. [62,63] Thereby, UV-Vis confirmed the final functionalization with the biologically relevant ligands, facilitating the nanoparticle interaction within the biological environment and their connection to ligand antigens. Besides inducing immunogenicity, the surface functionalization affected the nanoparticle dispersibility. In order to investigate the behaviour of the functionalized particles in solution, the particles’ hydrodynamic radius was examined using dynamic light scattering (DLS). As-functionalized nanoparticles showed good particle dispersion stability with a Z-Average of 59.9 nm (PDI: 0.25) for CA-capped particles and 73.0 nm (PDI: 0.16) for FA-capped particles (Figure 3E). When the particles were transferred from an initial DMSO solution into an aqueous phase to be applied to cells, the mean hydrodynamic radius decreased to 52.9 nm, proving the immunolabel’s suitability for biological investigations.

Taken together, these results indicated the successful functionalization of nanoparticles with folic and caffeic acid to bridge the particles’ spectral response to biological entities.

## 5 Single-Particle Cathodoluminescence and Energy Dispersive X-Ray Emission

To enable a reliable recognition of individual labels through ColorEM, it is imperative that the dopant-specific features remain detectable on a single-particle level. Making biological samples resistant to the vacuum conditions in TEM requires their fixation, dehydration and embedding in resin. [17] During each process, chemicals (e.g. buffer solutions, resin, heavy metals etc.) are used, which potentially veil or interfere with the label’s CL and/or EDX-SM signal. [20,64–66]

Initally, phase-transferred (BF_4_^–^-capped) particles deposited on a TEM grid investigated. In this set-up the interaction of the particles with their environment was kept to a minimum and non-radiative quenching effects reduced, highly favouring (CL)-emissive properties. These optimal imaging conditions allowed for benchmarking CL vs. EDX-SM signals at an early stage, outlined the methods potential. Figures 4A and B show the acquired CL response for single particles in scanning TEM (STEM), and the high-angle annular dark-field (HAADF) images indicate individual terbium- or europium-doped nanoparticles, which both exhibited a diameter of *≈*20 nm. For each pixel, the CL emission was simultaneously recorded to the HAADF image. The CL spectra were obtained by integrating an area on the particle and were compared with a similar-sized area on the carbon grid, which was in close proximity to the particle.

**Figure 4:**
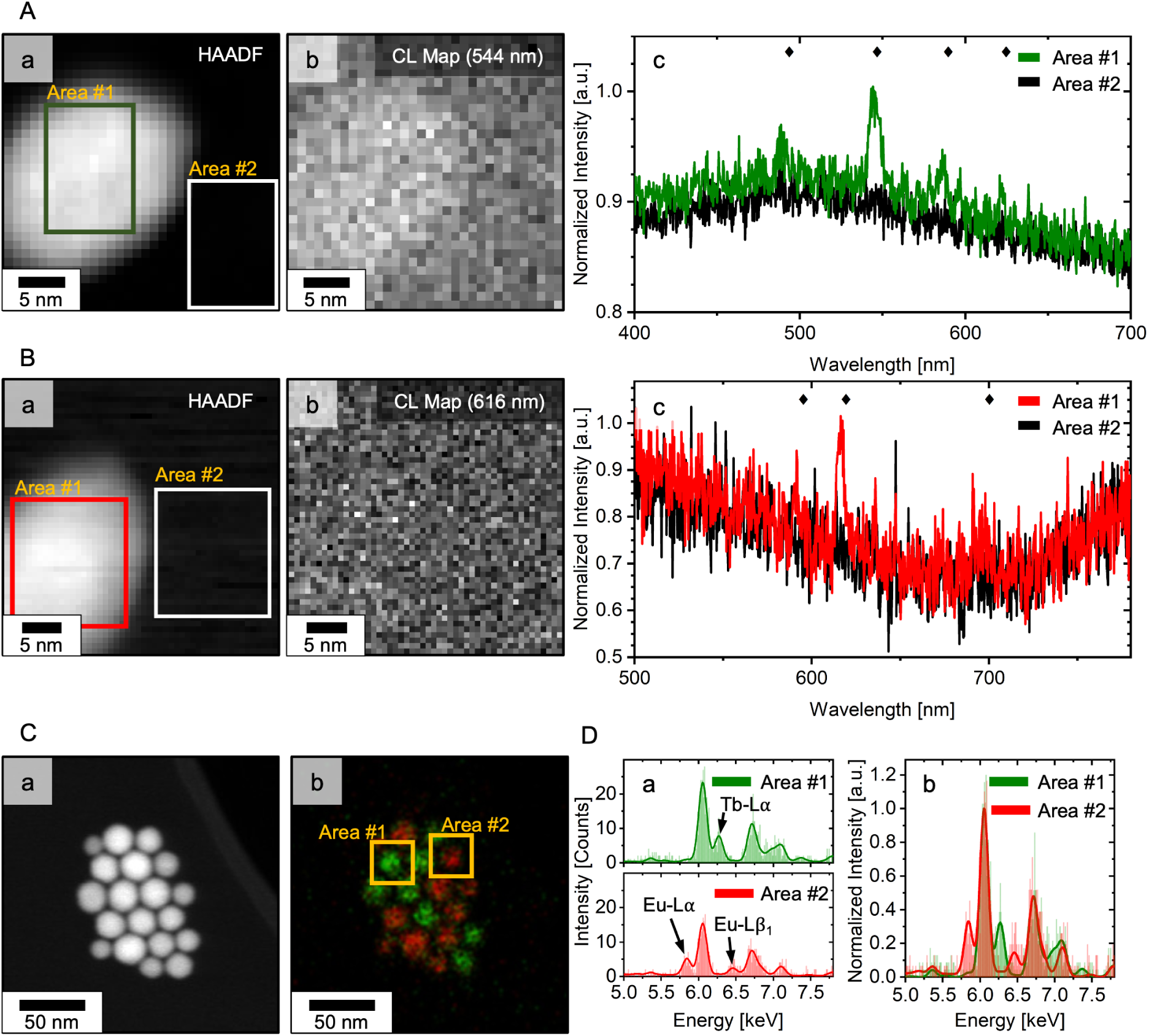
Single-Particle Recognition. (**A, B**) Single-particle CL, investigated on a TEM grid using a scanning TEM (STEM) set-up. High-angle annular dark-field (HAADF) images of terbium-doped (**A, a**) and europium-doped (**B, a**) core/shell nanoparticles acquired in parallel with the particles’ CL signal (**A, b; B, b**). Signal sliced on the main transitions (Tb: 544*±*14.3 nm; Eu 616*±*9.9 nm). (**A, c; B, c**) CL signal of the maps was integrated across the marked areas, #1 on the particle and #2 on the carbon film and corresponding spectra are shown. Area sizes: terbium-particles (green): 15x10 nm (15x10 px); europium-particles (red): 14.7x16.8 nm (24x21 px). (**C**) HAADF (**a**) and EDX-SM (**b**) images of mixed-doped particle ensemble (1:1, weight based) on a carbon grid. Coloration based on rare-earth separation. (**D, a**) EDX-SM spectra acquired from the identical ensemble and EDX-SM spectra derived from two identical areas (15.3x15.3 nm; 11x11 px in **C, b**), representing single particles. Characteristic lines labelled. (**D, b**) Normalized spectra of both areas a.u.: arbitrary units.

The CL spectrum of the terbium-doped particle (Figure 4A, c) was dominated by a peak at 544 nm. This was the most intense signal in the bulk spectrum and was previously assigned to the ^5^D_4_ *→*^7^ F_5_ transition of terbium. [41] Two additional emissions were found, in line with the ^5^D_4_ *→*^7^ F_4_(489 nm) and ^5^D_4_ *→*^7^ F_3_(586 nm) transitions of terbium. Thus, the emission spectrum of the small single particle matched that of the bulk sample, in which the three main transitions of terbium were recognized. Presuming the terbium dopant, the fourth transition (^5^D_4_ *→*^7^ F_3_; 489 nm) was identified, however, was not expressive enough and without data processing could not act as an identification feature. In contrast, the carbon-covering area did not exhibit any signal of terbium, demonstrating that STEM-CL did not give rise to non-local excitations, enabling a more precise correlation of position and signal. This demonstrated an advantage of STEM-CL over SEM-CL, as the high acceleration voltages (here *≥*60 kV) confine the lateral energy deposition and prevent vast excitations. [19,67] Moreover, the CL map, centered on the main transition at 544 nm, displayed the silhouette of the nanoparticle visible in the HAADF image with individual pixels on the particle giving an increased signal, while the carbon around the particle is unobtrusive. Thus, the location of the terbium particle is visible based on its CL response.

In contrast, the CL spectrum of the europium-doped nanoparticle (Figure 4B, c) showed one distinguishable signal without further data processing. It was located at 616 nm and matched the position of europium’s ^5^D_0_ *→*^7^ F_2_ transition, as previously observable in the bulk sample. Data processing (FFT filter, low pass) revealed further peaks at 590 nm and 691 nm, which were in harmony with the ^5^D_0_ *→*^7^ F_1_ and ^5^D_0_ *→*^7^ F_4_ transitions detected in the europium bulk sample (see Suppl. Figure S5). However, the poor data quality impeded a visualization of the particle within the transition sliced (616 nm) CL map.

Both CL maps covered an area of *≈*1,225 nm^2^, given pixel sizes of 1.0 nm(Tb) and 0.7 nm(Eu). Their total acquisition times were 307.2 s for the terbium- and 125 s for the europium-doped nanoparticles, whereby the particle spectra were derived from data recorded within 45 s and 25.2 s, respectively. These times were in similar orders of magnitude as those reported by Swearer *et al*. for their sub-20 nm cathodophores. In conclusion, the CL acquisition allowed the detection of dopant-specific ff-transition emissions for particles on a TEM grid and thus enabled their identification through CL-ColorEM.

Furthermore, the generated characteristic X-ray emission was investigated using a mixed-doped nanoparticle ensemble (1:1, weight based) deposited on a TEM grid (Figure 4C). The EDX-SM map closely resembled the HAADF image, whereby the individual particle outlines were recognizable based on the rare-earths’ characteristic X-rays. From the map, two areas were identified (Figure 4D), each area corresponding to the total signal generated by an individual particle (dimensions: 15.3x15.3 nm). The extracted spectra showed the Gd-Lα/β lines attributed to the host matrix, paired with the unique response of the rare-earth center. While area #1 exhibited a distinct emission at 6.27 keV, area #2 showed one at 5.84 keV. Using the previously determined key features, terbium and europium were assigned as dopants to area #1 and area #2, respectively (left signal-Eu, right signal-Tb). Beyond that, the energy resolution in EDX-SM enabled the distinction of neighboring atoms (Eu:63, Gd:64, Tb:65), which is an important asset for multi-color EDX-SM. The clear identification was accompanied by an excellent signal-to-noise ratio (SNR), already reached while scanning the depicted area (*≈*46,000 nm^2^; pixel size: 1.39 nm) for *≈*15 s.

Although both CL and EDX-SM showed the potential for single-particle recognition on a TEM grid-level, their signal quality differed. While the spectra of CL and EDX-SM were derived from similar-sized areas (real space), CL falls short compared to EDX-SM when assessing the SNR. Comparing the normalized spectra in Figure 4 A, c; 4B, c and 4D, b, it became evident that CL mainly suffered from a high background signal and readout noise, whereas the X-ray lines in EDX-SM emerged from the baseline and the elemental detection using the Velox software easily assigned the specific rare-earth. As shown in Suppl. Figure S5, data processing could reduce noise such that the dopant’s characteristic features were detectable, yet considering the acquisition parameters (e.g. mapped area, pixel size etc.) and especially the ’time-on-target’ needed in CL for sufficient SNR, EDX-SM proved to be the more potent recognition method for ColorEM. Taking the present acquisition times and considering the resulting data quality, even with a generous estimation CL would still require substantially longer mapping times (see Suppl. Figure S6). Beyond that, unlike EDX-SM, the CL acquisition required cooling to cryo temperatures in order to improve the SNR, but cannot be used reliably without data processing, resulting an increased effort in data acquisition and processing. Challenged by the fast acquisition speed and surprisingly good signal-to-noise ratio in EDX-SM, the limitations of reliable CL-based immunolabelling become evident. Despite the offered multi-color potential and successfully demonstrated single-particle recognition of sub-20 nm nanoparticles, the data quality (SNR, acquisition time etc.) did not meet the standards needed for the immunolabels application. While CL acquisition even of resin-embedded immunolabels in biological samples is conceptually feasible (see Suppl. Figure S7) and more thorough data processing might be implemented, the acquisition times and the SNR needed for a biologically relevant field-of-view (>100 µm^2^) lie beyond a currently achievable scope due to e.g. drift or beam damage of the labels. As the STEM used was equipped with a light detection device (Mönch, Attolight) fitted with a electron-multiplying CCDs (EMCCD), it ensured optimal photon detection. In terms of detection, the reduction of efficiency can be attributed to loss in the spectrometer (grating efficiency) where hardware advances hardly seem foreseeable, and the presence of background-related noise which is large with respect to the small cathodoluminescence signal of the labels. Brighter and more beam-stable labels are needed paired with adjustments in sample preparation (e.g. decrease section thickness, resin changes etc.) to utilize STEM-CL for immuolabelling. Furthermore, filters could be introduced and centered on a rare-earth line which coupled to a photon multiplier tube enable CL-immunolabelling. Although this would not reduce the background effect, it would essentially negate the reduction of signal related to the usage of spectrometers

In view of these results, where EDX-SM is at least on par with CL for its capability of multi-color immunolabelling, yet performs substantially better in signal generation and acquisition speed, holding onto CL did not appear justified; particularly with regard to upcoming emission-aggravating sample preparation steps and functionalizations.

## 6 Immunolabelling using ColorEM Label Differentiation

Given the superior qualities of EDX-SM in terms of SNR and collection times, its potential for immunolabelling was exploited. Folic acid-capped terbium-doped particles and caffeic acid-capped europium-doped nanoparticles were used to label biological structures (cell surface receptors) on the outer cell membrane. The labels’ attachment on the fixed cell surface and their different luminescent centers were investigated using a STEM-EDX-SM set-up (Figure 5). The cell’s ultrastructure, including different organelles such as the cell nucleus and mitochondria (Figure 5A) became accessible through to heavy metal staining (osmium, uranyl acetate and lead citrate). The nanoparticle labels were visible based on their strong Z-contrast and uniform size, and were found attached to the outer cell membrane at the microvilli of the cells, which are the most exposed and thus likeliest contact points for the nanoparticles (Figure 5A). No significant amounts of randomly scattered or agglomerated nanoparticles were found within the cells’ broader vicinity (Figure 5A, see also zoom-out in Figure 5E), suggesting the binding of the nanoparticle labels to structures on the cell membrane (see also bright-field TEM images in Suppl. Figure S8). The successful binding of folic acid-functionalized nanocrystals to folate receptors on the cell membrane was further supported by previous blocking experiments with free folic acid, showing greatly reduced receptor binding. [17] After sub-region selection that included multiple surface-immobilized nanoparticles (Figure 5B), EDX-SM mapping was performed to probe the labels’ nature (verification as label, dopant and related surface functionalization; Figure 5C, D). The EDX-SM map (Figure 5C) visualized both luminescent centers, which correlated well with the previously assigned particle positions. EDX-SM derived spectra of the following four representative areas were measured: (#1) europium-doped nanoparticle, (#2) terbium-doped nanoparticle, (#3) background on cell material, and (#4) background on epoxy resin (Figure 5D; locations depicted in Figure 5A, B). Using the characteristic X-ray pattern mainly based on the L*α* lines location, both dopants were unambiguously assigned to the particles and furthermore stood out well from the two background signals recorded at areas (iii) and (iv).

**Figure 5:**
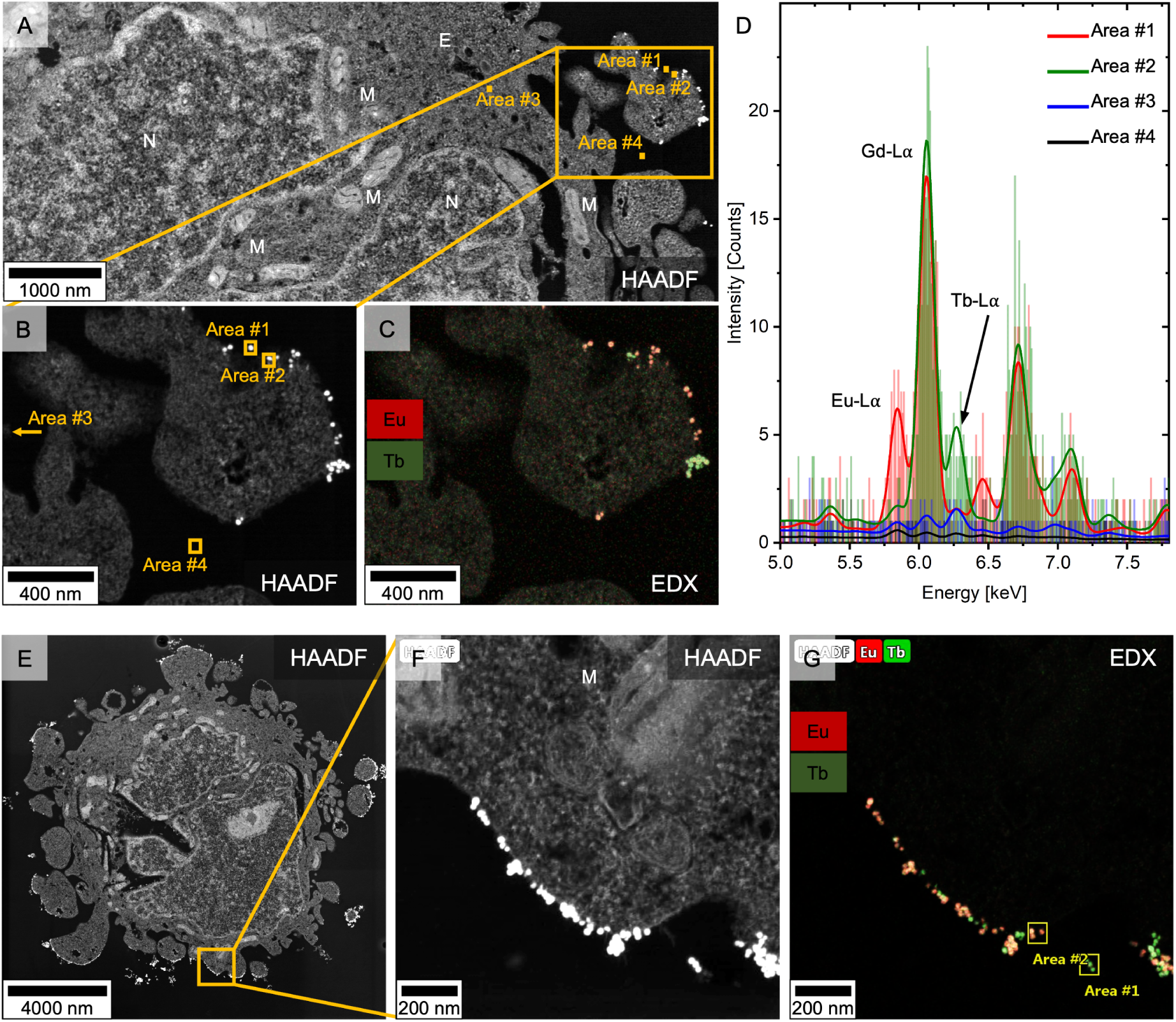
EDX-SM Immunolabelling. (**A**) HAADF overview image of epoxy-embedded HeLa cells after nanoparticle-based immunolabelling. Europium-doped nanoparticles capped with caffeic acid and terbium-doped nanoparticles capped with folic acid were used as labels. Cell components are indicated with E: endosome, M: mitochondria and N: nucleus. A selected subsection of the cell membrane at microvilli (**B**) was mapped using EDX-SM (**C**), and the spectra of four representative areas were extracted (**D**). The areas show exemplarily membrane-attached europium-(#1, red) and terbium-doped (#2, green) particles as well as the cell background (#3, blue), and cell- and particle-free epoxy resin background (#4, black). Also a second area of the same cell with multiple nanoparticles bound along the cell membrane closer to the cellular body than the microvilli investigated in **A-D** was measured using EDX-SM **(E-G)**.

Also a second area of the same cell (Figure 5E), where multiple nanoparticles were bound along the cell membrane closer to the cellular body than to the microvilli investigated in Figure 5A-D, was measured with EDX-SM (Figure 5E-G). The doping of the nanoparticles bound along the cell membrane as visualised in the high magnification HAADF image in Figure 5F was determined and depicted in the corresponding EDX-SM map (Figure 5G).

Strikingly, the EDX-SM map (Figure 5C) was recorded using a pixel size of 2.776 nm, which enabled the mapping of biologically relevant areas in a reasonable time frame while gaining insights in the cell’s ultrastructure. The acquisition parameters used resulted in a mapping speed of 109 s/µm^2^, which allowes up to whole-cell imaging at high resolutions, and provided an important asset to resolve structure-function relationships in detail. Notably, optimizing the acquisition parameters as well as the label’s composition and architecture could enhance mapping speed. Tailoring pixel size and dwell time towards a scientific question, redundant resolution can be traded for more area mapped during an equal timeframe. The speed of the dopant identification and thus the total mapping time is determined by the count rate generated by the immunolabel. As the characteristic lanthanide was located in the nanoparticle’s core and the passivating shell is presumably not a prerequisite for a strong EDX signal, smaller (sub-15 nm) yet brighter labels (higher local count rates) could be synthesized using pure NaEuF_4_ and NaTbF_4_ nanoparticles. Thereby, the immunolabelling precision would be improved, while the acquisition time would be reduced. [26]

For the immunolabelling approach presented here, EDX-SM delivered on its promise by providing a fast and reliable tool for label localization. In addition to achieving a well preserved and visible ultrastructure with this protocol, each laboratory-designed label could be classified based on its luminescent center and enabled the localization of the cellular components *via* its surface functionalization. The introduced labels were easily retrieved, as interfering electron-dense structures (e.g. osmium precipitates) were eliminated from consideration owing to their X-ray signature. The lanthanide centers provided an inherent background-free option, as their characteristic X-rays were not obscured by elements found naturally in biological systems (e.g. C, N, S, etc.) or added during sample preparation (e.g. resin, heavy metals).

In contrast, upon electron irradiation, epoxy resins are known to produce auto-luminescence in the visible range, which impedes the lanthanides’ CL emissions (see Suppl. Figure S7). [20] Moreover, EDX detectors are offered in many STEM systems, which allows more widespread access and use of EDX-based ColorEM compared to CL that additionally also requires cryo temperatures during data acquisition and often more data processing. Through lanthanide substitution, the library of the immunolabels presented here is expandable, allowing multiple colours with unique (X-ray fingerprints to be used for the elucidation of structure-function relationships at the nanometer scale.

## 7 Conclusion

In this work, we synthesized sub-20 nm nanoparticles as immunolabels and offered the first practical method for multi-colour labelling of biological structures in (STEM)-ColorEM. The core/shell nanoparticles we presented shared a similar size, shape and composition of the shell’s material, but differed fundamentally in their core’s elemental composition. Europium and terbium were incorporated in a sodium gadolinium fluoride host matrix acting as luminescent centers. They enabled single-particle recognition based on their unique spectral properties (CL) and characteristic X-ray signature (EDX). Each dopant was given a surface functionalization, aiming to target cell surface receptors in an immunolabelling approach. The folic acid and caffeic acid-capped particles were rapidly (co)-localized and identified using EDX-SM and simultaneously contextualized in the cell’s ultrastructure obtained by HAADF imaging. With EDX-SM and our highly-doped immunolabels, we overcame the limiting slow acquisition speeds which historically hampered the use of EDX-SM in the biological field, and reached a fast mapping of biologically relevant areas within seconds to minutes, while offering 3 nm resolution. Tailoring the immunolabels towards EDX-ColorEM by increasing the characteristic lanthanide content while reducing their size holds the promise of further improving acquisition time and localization precision. Exchanging the doped lanthanide atoms offers the potential to expand the library of presented immunolabels without interfering with each other or with cell ultrastructure staining, out-competing standard gold immunolabeling procedures that are purely based on Z-contrast. We showed that CL failed the translation towards application, as it required significantly longer acquisition times (>100 times) compared to EDX-SM and delivered a inferior signal-to-noise ratio. In order to successfully transfer CL into ColorEM application, brighter and more beam-stable CL labels are required. Additionally, novel approaches involving filters and photon counting devices could reduce the background signal and applied alongside adjustments in sample preparations. However, competing against EDX-recognition in ColorEM with EDX detectors being widely implemented in various STEM systems, and less effort required for data acquisition (cooling, data processing etc.), the question remains whether CL provides the best method to image epoxy-embedded samples.

## 8 Materials and Methods

### 8.1 Chemicals

All materials were purchased from Sigma-Aldrich and used without further purification if not stated otherwise. Oleic acid (technical grade, 90%), 1-Octadecene (technical grade, 90%), 2-Aminoethyl-dihydrogen-phosphate (AEP), sodium hydroxide (NaOH), nitrosyltetrafluoroborate (95%), ammonium fluoride (puriss, p.a., ACS reagent, *≥*98%), chloroform (ReagentPlus, *≥*99.8%, 0.5-1.0% ethanol as stabilizer), dimethylsulfoxide (DMSO, anhydrous, *≥*99.9%), folic acid (FA, *≥*97%), caffeic acid (CA, HPLC, *≥*98.0%), cyclohexane (ROTISOLV^®^, HPLC), ethanol (absolute, ACS reagent, *≥*99.8%, Honeywell), dimethylformamide (DMF, ReagentPlus, *≥*99%), methanol (Fisher Scientific, HPLC Gradient grade, *≥*99.9%), *N* -Hydroxysuccinimide (98%), sodium trifluoroacetate (98%), europium(III)-acetate hydrate (99.9% trace metals basis), gadolinium(III)-acetate hydrate (99.9% trace metals basis), terbium(III)-acetate hydrate (99.9% trace metals basis), 1-ethyl-3-(3-dimethylaminopropyl)carbodiimid (EDC), glutaraldehyde solution (GA, Grade I, 50% in H_2_O, specially purified for use as an electron microscopy fixative or other sophisticated use), epoxy embedding kit 45359, 4% PFA solution (prepared from solid; paraformaldehyde, meets analytical specification of DAC, 95%-100.5%), ultrapure hydrogen peroxide (Suprapur, Merck, Germany), ultrapure nitric acid (67%, Normatom, VWR Chemicals). Osmium tetroxide solution (4%) and sodium cacodylate buffer (0.2M, pH 7.4) were purchased from Electron Microscopy Sciences (EMS). Dulbecco’s Phosphate Buffered Saline (PBS, Modified, without calcium chloride and magnesium chloride, liquid, sterile filtered, suitable for cell culture), folic acid-free medium (prepared from: Dulbecco’s Modified Eagle’s Medium - low glucose, 10 x, with 1000 mg/L glucose (1x), without L-glutamine, sodium bicarbonate, and folic acid, liquid, sterile-filtered, suitable for cell culture)

### 8.2 Preparation of Core/Shell Particles

#### 8.2.1 Preparation of Core Particles

A reaction protocol according to Wang *et al.*[68] was used. Briefly, rare-earth acetates (a total amount of 1 mmol) were added in the desired stoichiometry to a mixture consisting of 10 mL oleic acid and 15 mL octadecene. The solution was degassed at room temperature under vacuum, and the temperature was gradually increased to 120 *^◦^*C and kept at this temperature for 90 min under vacuum and vigorous stirring. After cooling to room temperature, 2.5 mL of methanolic NaOH solution (1 M) and 7.75 mL of methanolic NH_4_F solution (0.4 M) were added under nitrogen to the reaction solution. The solution was slowly exposed to a vacuum, and the methanol was removed from the reaction, first at room temperature, and then the temperature was subsequently increased to 50 *^◦^*C and kept under vacuum for 30 min. The temperature was increased to 100 *^◦^*C and cycled three times between vacuum and nitrogen atmosphere. The solution remained under a nitrogen atmosphere, and the temperature was increased to approximately 300 *^◦^*C within 30 min. The reaction remained at this temperature for 1 h before the heating mantle was removed and the reaction was cooled down to room temperature. The yellowish turbid solution was transferred into falcon tubes, and the particles were precipitated by adding ethanol (approximately 10 mL). The particles were separated from the solution by centrifugation (15 min, 6000 rcf, 20 *^◦^*C), the supernatant was decanted, and the precipitate was collected in clean cyclohexane. After washing three times, the clean particles were centrifuged at low speed (20 min, 200 rcf, 20 *^◦^*C) without precipitation to remove big agglomerates and precipitated sodium fluoride. The solution was syringe filtered (Minisart SRP, hydrophobic, PTFE, 25 mm, 0.2 µm) and stored in a glass vial at, 4 *^◦^*C under exclusion of light.

#### 8.2.2 Shelling Procedure (Growth/Surface Passivation)

The nanoparticles’ diameter was first increased to enlarge the luminescent core, which was then passivated with a non-doped shell using a hot-injection protocol based on Siefe *et al.*[69] This process was divided in (i) the preparation of precursor solutions and (ii) the time-dependent feeding of precursor solutions using a syringe pump.

- Rare-earth precursor (0.1 M): For a typical reaction a total amount of 2 mmol rare-earth acetate (composition dependent on growth or passivation step) was added in a solution consisting of 8 mL oleic acid and 12 mL octadecene. The solution was degassed at room temperature under vacuum, and the temperature was gradually increased to 140 *^◦^*C and kept at this temperature for 60 min under vacuum and vigorous stirring. Then the solution was cooled to room temperature and transferred to a syringe protected by nitrogen or argon atmosphere.
- Sodium trifluoroacetate (0.4 M): 3.2 mmol of Na-TFA was added to 8 mL oleic acid and treated as above.

During each hot injection step, *≈*100 mg of the previous particles were used. Thereby, the core particles usually exhibited a diameter of d_TEM_*≈*9-10 nm and the grown particles a d_TEM_*≈*13-14 nm. The particles were added in a solution of 4 mL oleic acid and 8 mL octadecene. Cyclohexane was removed by keeping the reaction at 70 *^◦^*C for 30 min under vacuum. After increasing the temperature to 100 *^◦^*C and cycling the nitrogen atmosphere and vacuum three times, the nitrogen atmosphere was retained, and the temperature was ramped up to 300 *^◦^*C within 30 min. The precursors were added separately to the solution in time-separated cycles using syringe pumps (NE-1000, KFTechnology). The feeding parameters are shown in table 1. After completion of the last cycle, the solution was kept at 300 *^◦^*C for further 30 min and was then cooled down to room temperature.

**Table 1:** Parameters of the shelling process.

|  | Cycle 1-9 | Feeding<br>Rate | Total<br>Volume | Total Amount<br>of Substance |
| --- | --- | --- | --- | --- |
| RE Precursor | 1.86 mL | $0.3 \text{ mL min}^{-1}$ | 16.74 mL | 1.674 mol |
| Na-TFA | 0.465 mL | $0.303 \text{ mL min}^{-1}$ | 4.185 mL | 1.674 mol |

The yellowish turbid solution was transferred into falcon tubes, and the particles were precipitated by adding ethanol (approximately 10 mL). Then, the particles were separated from the solution by centrifugation (15 min, 6000 rcf, 20 *^◦^*C), the supernatant was decanted, and the precipitate was collected in clean cyclohexane. After washing three times, the colorless particle solution was centrifuged without precipitation at low speed (20 min, 200 rcf, 20 *^◦^*C) to remove big agglomerates, filtered using a syringe filter (Minisart SRP, hydrophobic, PTFE, 25 mm, 0.2 µm) and stored in a glass vial at 4 *^◦^*C under exclusion of light.

### 8.3 Functionalizations

#### 8.3.1 Phase-Transfer of Oleic Acid-capped Particles

The particles were phase-transferred into a polar medium according to a protocol of Dong *et al.*[51] Briefly, nitrosyl tetrafluoroborate (NOBF_4_, 100 mg) was dissolved in 3 mL DMF and layered with the same amount of oleic acid-capped particles dissolved in 3 mL cyclohexane. The two-phase system was then stirred at room temperature for 1 h. The cyclohexane layer was discarded, and the polar phase was transferred to a falcon tube. Chloroform (20 mL) was added, and the precipitated particles were collected by centrifugation (45 min, 9000 rcf, 20 *^◦^*C). After repeated purification with DMF/chloroform, the particles were redispersed in DMF and stored in a glass vial at 4 *^◦^*C under the exclusion of light.

#### 8.3.2 Ligand Exchange with 2-Aminoethyl Dihydrogen Phosphate (AEP)

The AEP-functionalization was carried out using an adapted protocol of Liu *et al.*[54] Specifically, AEP (200 mg) was dissolved under ultrasonic conditions in 15 mL milli-Q water and ethanol (3:2 v/v) solution. Particles dissolved in DMF were added to the solution and stirred for 48 h at room temperature. The particles were purified using the milli-Q water and ethanol (3:2 v/v) solution and collected by centrifugation (35 min, 11000 rcf, 15 *^◦^*C). The particles were redispersed in water, and the opalescent solution was stored after repeated washing cycles and syringe filtering at room temperature.

#### 8.3.3 Folic Acid- and Caffeic Acid-Coordination on AEP-capped Particles

The AEP-coordinated particles were functionalized with folic acid using a common EDC/NHS procedure. Accordingly, folic acid (12.0 mg, 0.027 mmol) and EDC (4.0 mg, 0.025 mmol) both were dissolved in 0.5 mL DMSO under ultrasonic conditions and merged together. NHS (13.42 mg, 0.116 mmol) was dissolved in 0.5 mL milli-Q water and added in the folic acid-containing solution. The solution was stirred for 1 h under the exclusion of light. The AEP-capped particles (70 mg in MES-Buffer) were added in the solution and stirred over night under the exclusion of light. The particles were purified by centrifugation (30 min, 11000 rcf, 15 *^◦^*C) using a 50 vol.% DMSO/water solution. The purification was completed when the supernatant of the solution after centrifugation (15 min, 6000 rcf, 15 *^◦^*C) showed no coloration by excess folic acid. The orange particles were taken up in DMSO and stored under the exclusion of light at 4 *^◦^*C. For the caffeic acid-capped particles, folic acid was substituted, and caffeic acid (14.59 mg, 0.081 mmol), EDC (13.97 mg, 0.090 mmol) and NHS (20.71 mg, 0.180 mmol) used during the reaction (for 100 mg AEP-capped particles).

#### 8.3.4 Cell Fixation and Immunolabeling Procedure

HeLa cells (ATCC CRM-CCL-2) were used for the labelling, as they are known for folate receptor overexpression and carry β-adrenergic receptors.[17,70] Further, they carry TGF-β_1_ receptors. [71] Cells were cultivated in Minimum Essential Medium Eagle (M2279, Sigma-Aldrich) supplemented with 10% fetal calf serum (FCS), and 1% of l-glutamine, penicillin–streptomycin, non-essential amino acids and sodium pyruvate at 37 *^◦^*C in 5% CO_2_ atmosphere. For labelling experiments, first, subconfluent cells were detached from a T75 culture flask using trypsin. Trypzination was stopped using folic acid-free FCS containing cell medium and cells were then centrifuged (5 min, 200 g), redispersed in fresh PBS buffer and counted using a Neubauer chamber. For each experimental condition, *≈*450 thousand cells were transferred to 1.5 mL Eppendorf tubes, centrifuged again, and then re-suspended and fixed in 4%PFA/1%GA for 2 h at room temperature. After an initial wash with PBS (750 µL), the cells were washed twice using 0.9% NaCl solution prior to incubation with the particle solution (10% DMSO-containing particle solution: 100 µg mL*^−^*^1^; 1000 µL) for 2 h at room temperature. Excess particles were removed by centrifugation (5 min, 300 g) and the cells additionally washed using 0.9% NaCl solution (750 µL). Lastly, NaCl solution was replaced by Na-cacodylate (2% GA, 0.1 M, pH 7.4, 500 µL) buffer solution and cells were stored at 4 *^◦^*C under the exclusion of light until sample preparation for electron microscopy (less than a week).

### 8.4 Characterization methods

#### 8.4.1 Electron Microscopy (TEM, STEM, EDX-SM, HAADF)

For transmission electron microscopy (TEM), scanning TEM (STEM), energy-dispersive X-ray spectromicroscopy (EDX-SM) and high-angle annular dark-field (HAADF) imaging of the nanoparticles without cells, the nanoparticle samples were prepared on holey carbon film-coated copper grids (200 mesh, EM Resolutions). Specifically, 2 µL of the purified particle stock solutions (*≈* 50 mg mL*^−^*^1^) were transferred to a separate vial and 1000 µL clean cyclohexane was added. The solution was homogenized under ultrasonic conditions for 5 min and 15 µL of the diluted solution was applied to the grids, then the cyclohexane was evaporated. Shortly before the measurement, the samples were cleaned in bubbles of ethanol over activated carbon to remove organic residues, which lead to C-buildup during STEM-EDX analysis.[72] To image the morphology and development stages of europium and terbium-doped nanoparticles with TEM, a Zeiss EM900 transmission electron microscope was used operating at 80 kV. The elemental composition of the particles was studied with STEM-EDX-SM and HAADF using a Talos F200X microscope at an accelerating voltage of 200 kV. There, the samples were mounted on a double-tilt holder and fixed using a copper ring and clamp. The data was processed using the software Velox 3.0.0.815 (FEI, USA) and Fiji. The size distribution was automatically determined using the ParticleSizer module in Fiji.[73] At least 2680 particles were measured.

For TEM, STEM-EDX-SM and HAADF imaging of HeLa cells labeled at their outer membrane with europium-doped nanoparticles capped with caffeic acid or terbium-doped nanoparticles capped with folic acid, the HeLa cells were treated as described above. After storage in Na-cacodylate buffer, the cell samples in 1.5 mL Eppendorf tubes were post-fixed and stained with 1 % osmium tetroxide in 0.1 M Na-cacodylate buffer for 1 h in the dark, followed by three washes with MilliQ water for 3 min each. The samples were then dehydrated with an Ethanol series (30 %, 50 %, 70 %, 90 % for 5 min each, 3 x 10 min 100 % Ethanol) and incubated for 1 h in a 1:1 mixture of 100 % Ethanol and 100 % Epon 812 substitute (Epoxy embedding kit 45359, Sigma-Aldrich). After an overnight incubation with 100 % Epon in 1.5 mL Eppendorf tubes with open lids, the cell samples were embedded in fresh 100 % Epon in the same tubes for at least 48 h at 60 *^◦^*C. After excising the Epon blocks from the Eppendorf tubes, ultrathin sections of 80-100 nm were cut with an ultramicrotome (Leica EM UC6) and applied onto Formvar-coated copper grids (100 mesh, EM Resolutions) for quality assessment with the Zeiss EM900 TEM, and onto Formvar-carboncoated copper grids (100 mesh, EM Resolutions) for STEM CL/EDX-SM/HAADF imaging with the Talos F200X microscope or Attolight Allalin electron microscope (CL bulk). Before imaging, the sections were contrasted for 20 min with 2 % uranyl acetate followed by 1 min of lead citrate; except for CL analysis, shown in **??**. The CL investigations were carried out using Chromatem (see CL methods section).

#### 8.4.2 Dynamic Light Scattering (DLS)

The hydrodynamic radius of the particles was determined by dynamic light scattering measurements at a Zetasizer Nano ZS90 instrument (Malvern Instruments Ltd., Worcestershire, United Kingdom) with 90*^◦^* scattering angle. Dependent on the functionalization step, the particles were present in cyclohexane, DMF, water and DMSO. The samples in cyclohexane and DMF were measured using a 3500 µL quartz cuvette (Thorlabs, CV10Q35A) and the samples in water and DMSO were measured in single-use cuvettes. The corresponding viscosities and refractive indices were taken from the instrument-internal database according to the solvent and sample. Typical sample preparation involved direct measurement of the particles in the stock solution. The attenuator was kept between 7-9, and a range between 200 kcps and 400 kcps was provided. For the measurements, three runs of 15 individual measurements were performed.

#### 8.4.3 Fourier-Transform Infrared Spectroscopy (FTIR)

Infrared spectroscopy was performed using a Bruker Tensor 27 instrument (Bruker Optics, Ettlingen, Germany) with a single reflection attenuated total reflectance (GladiATR) accessory (Pike Technologies, Fitchburg, Wisconsin, United States). The instrument was first filled with liquid nitrogen 30 min before the measurement. Typically, 32 scans were made in the range between 4000 cm*^−^*^1^ and 600 cm*^−^*^1^ with a resolution of 4 cm*^−^*^1^. The transmittance of the background was determined and subtracted from the sample.

#### 8.4.4 Inductively coupled plasma optical emission spectrometry (ICP-OES)

For composition analysis of the particles using inductively coupled plasma optical emission spectrometry (ICP-OES), 1-2 mg of the particles, dried from cyclohexane, were transferred to a quartz test tube. For digestion 300 µL ultrapure nitric acid (67%, Normatom, VWR Chemicals) and 300 µL ultrapure hydrogen peroxide (Suprapur, Merck, Germany) were added to the quartz test tube. The samples were digested at 1000-1200 W, 200 *^◦^*C and 120 bar for 10 min using a microwave (turboWAVE Inert, MWS GmbH). The digested solutions were transferred to previously weighted falcon tubes (15 mL) and diluted with ultrapure water to *≈*10 mL. The falcon tubes were weighed again for a precise determination of the total volume. The samples were then measured using an Agilent 5110 ICP-OES instrument calibrated using external standards for inorganic elements such as boron, europium, gadolinium, sodium and terbium. The system was calibrated in a range of 0-5 ppm and the elements measured in 2% HNO_3_ solution.

#### 8.4.5 Cathodoluminescence (CL)

The bulk CL spectra were measured using an Attolight Allalin electron microscope operated at 3 kV and 120 pA probe current. A 300 l/mm grating was used to optimize the spectrum’s resolution, and the spectrometer slits closed to 10 µm. In order to minimize beam damage, a 10 µm FOV was scanned fast. The measurements were performed *in situ* using a Nd:YAG reference sample to correct for slight wavelengths inaccuracies. The centre wavelength accuracy was *<*0.1 nm. The STEM-CL set was a custom-modified NION HERMES 200, fitted with a Mönch Attolight cathodoluminescence system, denoted as ’chromatem’. The data was recorded using the following acquisition parameters: (i) terbium-doped particle: spectral image 32x32 px, 300 ms per pixel dwell time, EMCCD mode x950, grating 150 gmm, blazed at 500 nm and centred at 500 nm, 350 pA, 10 mrd half angle. (ii) europium-doped particle: spectral image 50x50 px, 50 ms per pixel dwell time, EMCCD mode x950, grating 150 gmm, blazed at 500 nm and centred at 500 nm, 350 pA, 10 mrd half angle.

#### 8.4.6 Photoluminescence (PL)

Photoluminescence excitation and emission spectra were acquired using Horiba Fluoromax Plus Spectrometer using an excitation wavelength of λ_ex_= 395 nm for europium with 2.5 nm, 2.5 nm excitation and 2.5 nm, 2.5 nm emission slits. The measurement parameters for the terbium nanoparticles were the following: λ_ex_= 369 nm; Excitation slits: 5 nm, 5 nm; Emission slits: 2 nm, 2 nm; Grating density 1200 (Blaze 500). Measurements were conducted using 3500 µL quartz cuvettes (Thorlabs, CV10Q35A) and recorded in a range of 400(Eu: 450)-750 nm.

#### 8.4.7 Thermogravimetric Analysis (TGA)

The samples were dried in a vacuum oven at 40 *^◦^*C, and the particles were transferred into a 85 µL Al_3_O_3_ crucible. The measurement was conducted using a NETZSCH TG 209 F1 instrument (NETZSCH-Geraetebau GmbH, Selb, Germany), and the samples were heated under a nitrogen atmosphere from 25 *^◦^*C to 900 *^◦^*C at a heating rate of 10 *^◦^*C min*^−^*^1^. A typical mass of 3 mg of 5 mg was used per measurement.

#### 8.4.8 Ultraviolet-Visible (UV-Vis) Spectroscopy

The absorption spectra of the particles were recorded using a Jenway 6705 UV/Vis spectrometer. All data lines were corrected for solvent absorbance by a baseline. Concentrations were adjusted not to exceed a maximum absorbance of 3 arbitrary units (a.u). Samples were measured using SARSTEDT Polystyrol/ Polystyrene 3.5 ml cuvettes.

## Acknowledgements

We would like to express our sincere gratitude to Dr. Kerda Keevend for her contributions to this research project and previous research in the field of cathodoluminescent nanoparticles. We further would like to thank the ScopeM imaging center of ETH Zurich for access to their EDX-SM electron microscopes and their technical support. This project has been funded in part by the Swiss National Science Foundation (SNSF, Eccellenza grant no. 181290), the Swiss Cancer Research Foundation (KFS-4868-08-2019), the National Agency for Research under the program of future investment TEMPOS-CHROMATEM (reference no. ANR-10-EQPX-50)and European Union’s Horizon 2020 research and innovation programme under Grant Agreements 823717 (ESTEEM3).

## Author Contributions

S.H. performed main experiments (e.g nanoparticle synthesis and functionalization), analyzed their data and drafted the manuscript. L.R.H.G. helped with the development of the immunolabelling protocol and cell cultivation. M.K. performed single-particle CL investigations. C.M. performed CL bulk assessment of samples. V.K. supervised the sample preparation for electron microscopy, performed post-staining of the sections and analysed the ultrastructure of the cell. A.G. developed and performed elemental analysis protocols, conducted EDX-SM analysis of particles and sections and provided guidance throughout the study. I.K.H conceived and supervised the study and edited the manuscript. All authors contributed to the manuscript writing and have given approval to the final version of the manuscript.

## References

[1] B. Titze, C. Genoud, Biology of the cell 2016, 108, 307–323.

[2] S. W. Hell, Science 2007, 316, 1153–1158.

[3] T. A. Klar, S. W. Hell, Opt. Lett. 1999, 24, 954–956.

[4] S. W. Hell, S. J. Sahl, M. Bates, X. Zhuang, R. Heintzmann, M. J. Booth, J. Bewersdorf, G. Shtengel, H. Hess, P. Tinnefeld, A. Honigmann, S. Jakobs, I. Testa, L. Cognet, B. Lounis, H. Ewers, S. J. Davis, C. Eggeling, D. Klenerman, K. I. Willig, G. Vicidomini, M. Castello, A. Diaspro, T. Cordes, J. Phys. D: Appl. Phys. 2015, 48, 443001.

[5] P. de Boer, J. P. Hoogenboom, B. N. G. Giepmans, Nat. Methods 2015, 12, 503–513.

[6] G. E. Palade, J. Biophys. Biochem. Cytol. 1955, 1, 59–68.

[7] T. Svitkina in Cytoskeleton Methods and Protocols: Methods and Protocols, (Ed.: R. H. Gavin), Methods in Molecular Biology, Springer, New York, NY, 2016, pp. 99–118.

[8] D. P. Hoffman, G. Shtengel, C. S. Xu, K. R. Campbell, M. Freeman, L. Wang, D. E. Milkie, H. A. Pasolli, N. Iyer, J. A. Bogovic, D. R. Stabley, A. Shirinifard, S. Pang, D. Peale, K. Schaefer, W. Pomp, C.-L. Chang, J. Lippincott-Schwartz, T. Kirchhausen, D. J. Solecki, E. Betzig, H. F. Hess, Science 2020, 367, eaaz5357.

[9] N. M. Pirozzi, J. P. Hoogenboom, B. N. G. Giepmans, Histochem. Cell Biol. 2018, 150, 509–520.

[10] K. Keevend, L. Puust, K. Kurvits, L. R. H. Gerken, F. H. L. Starsich, J.-H. Li, M. T. Matter, A. Spyrogianni, G. A. Sotiriou, M. Stiefel, I. K. Herrmann, Nano Lett. 2019, 19, 6013–6018.

[11] B. Gu, Q. Zhang, Adv. Sci. 2018, 5, 1700609.

[12] X. Zhu, J. Zhang, J. Liu, Y. Zhang, Adv. Sci. 2019, 6, 1901358.

[13] B. Zheng, J. Fan, B. Chen, X. Qin, J. Wang, F. Wang, R. Deng, X. Liu, Chem. Rev. 2022, 122, 5519–5603.

[14] M. B. Prigozhin, P. C. Maurer, A. M. Courtis, N. Liu, M. D. Wisser, C. Siefe, B. Tian, E. Chan, G. Song, S. Fischer, S. Aloni, D. F. Ogletree, E. S. Barnard, L.-M. Joubert, J. Rao, A. P. Alivisatos, R. M. Macfarlane, B. E. Cohen, Y. Cui, J. A. Dionne, S. Chu, Nat. Nanotechnol. 2019, 14, 420–425.

[15] M. Banski, A. Podhorodecki, J. Misiewicz, M. Afzaal, A. L. Abdelhady, P. O’Brien, J. Mater. Chem. C 2013, 1, 801–807.

[16] K. Keevend, T. Coenen, I. K. Herrmann, Nanoscale 2020, 12, 15588–15603.

[17] K. Keevend, R. Krummenacher, E. Kungas, L. R. H. Gerken, A. Gogos, M. Stiefel, I. K. Herrmann, Small 2020, 16, 2004615.

[18] D. F. Swearer, S. Fischer, D. K. Angell, C. Siefe, A. P. Alivisatos, S. Chu, J. A. Dionne, ACS Photonics 2021, 8, 1539–1547.

[19] S. A. Rehman, J. B. Conway, A. Nichols, E. R. Soucy, A. Dee, K. Stevens, S. Merminod, I. MacNaughton, A. Curtis, M. B. Prigozhin, Lanthanide Cathodophores for Multicolor Electron Microscopy, 2023.

[20] A. Srinivasa Raja, P. de Boer, B. N. G. Giepmans, J. P. Hoogenboom, Macromol. Biosci. 2021, 21, 2100192.

[21] M. Scotuzzi, J. Kuipers, D. I. Wensveen, P. de Boer, K. (W. Hagen, J. P. Hoogenboom, B. N. G. Giepmans, Sci. Rep. 2017, 7, 45970.

[22] B. N. G. Giepmans, T. J. Deerinck, B. L. Smarr, Y. Z. Jones, M. H. Ellisman, Nat. Methods 2005, 2, 743–749.

[23] H. Herd, N. Daum, A. T. Jones, H. Huwer, H. Ghandehari, C.-M. Lehr, ACS Nano 2013, 7, 1961–1973.

[24] M. F. Hoes, N. Grote Beverborg, J. D. Kijlstra, J. Kuipers, D. W. Swinkels, B. N. Giepmans, R. J. Rodenburg, D. J. van Veldhuisen, R. A. de Boer, P. van der Meer, Eur. J. Heart Fail. 2018, 20, 910–919.

[25] P. de Boer, N. M. Pirozzi, A. H. G. Wolters, J. Kuipers, I. Kusmartseva, M. A. Atkinson, M. Campbell-Thompson, B. N. G. Giepmans, Nat. Commun. 2020, 11, 2475.

[26] M. Amiry-Moghaddam, O. P. Ottersen, Nat. Neurosci. 2013, 16, 798–804.

[27] S. Wen, J. Zhou, K. Zheng, A. Bednarkiewicz, X. Liu, D. Jin, Nat. Commun. 2018, 9, 2415.

[28] N. J. J. Johnson, S. He, S. Diao, E. M. Chan, H. Dai, A. Almutairi, J. Am. Chem. Soc. 2017, 139, 3275–3282.

[29] Q. Su, S. Han, X. Xie, H. Zhu, H. Chen, C.-K. Chen, R.-S. Liu, X. Chen, F. Wang, X. Liu, J. Am. Chem. Soc. 2012, 134, 20849–20857.

[30] Y. Lu, E. Sega, C. P. Leamon, P. S. Low, Adv. Drug Deliver. Rev., Folate Receptor-Targeted Drugs for Cancer and Inflammatory Diseases 2004, 56, 1161–1176.

[31] A. Bosak, A. Knežević, I. Gazić Smilović, G. Šinko, Z. Kovarik, J. Enzyme Inhib. Med. Chem. 2017, 32, 789–797.

[32] J. F. Hainfeld, F. R. Furuya, J. Histochem. Cytochem. 1992, 40, 177–184.

[33] J. Gu, M. D’Andrea, Am. J. Anat. 1989, 185, 264–270.

[34] S. Yokota, J. Histochem. Cytochem. 1988, 36, 107–109.

[35] M. Howarth, W. Liu, S. Puthenveetil, Y. Zheng, L. F. Marshall, M. M. Schmidt, K. D. Wittrup, M. G. Bawendi, A. Y. Ting, Nat. Methods 2008, 5, 397–399.

[36] B. D. Chithrani, A. A. Ghazani, W. C. W. Chan, Nano Lett. 2006, 6, 662–668.

[37] M. A. Islam, S. Barua, D. Barua, BMC Syst. Biol. 2017, 11, 113.

[38] G. Bao, K.-L. Wong, D. Jin, P. A. Tanner, Light Sci. Appl. 2018, 7, 96.

[39] K. Binnemans, Coordin. Chem. Rev. 2015, 295, 1–45.

[40] Z. Boruc, B. Fetlinski, M. Kaczkan, S. Turczynski, D. Pawlak, M. Malinowski, J. Alloy. Compd. 2012, 532, 92–97.

[41] K. Keevend, M. Stiefel, A. L. Neuer, M. T. Matter, A. Neels, S. Bertazzo, I. K. Herrmann, Nanoscale 2017, 9, 4383–4387.

[42] J. A. Bearden, Rev. Mod. Phys. 1967, 39, 78–124.

[43] C. Chen, J. Ke, X. E. Zhou, W. Yi, J. S. Brunzelle, J. Li, E.-L. Yong, H. E. Xu, K. Melcher, Nature 2013, 500, 486–489.

[44] H. Silva, N. M. F. Lopes, Front. Physiol. 2020, 11.

[45] C. D. Strader, M. R. Candelore, W. S. Hill, I. S. Sigal, R. A. Dixon, J. Biol. Chem. 1989, 264, 13572–13578.

[46] G. Swaminath, X. Deupi, T. W. Lee, W. Zhu, F. S. Thian, T. S. Kobilka, B. Kobilka, J. Biol. Chem. 2005, 280, 22165–22171.

[47] C.-W. Huang, S.-Y. Lee, T.-T. Wei, Y.-H. Kuo, S.-T. Wu, H.-C. Ku, Biomed. Pharmacother. 2021, 142, 112028.

[48] T. Cao, T. Yang, Y. Gao, Y. Yang, H. Hu, F. Li, Inorg. Chem. Commun. 2010, 13, 392– 394.

[49] K. Yang, H. Peng, Y. Wen, N. Li, Appl. Surf. Sci. 2010, 256, 3093–3097.

[50] V. Venkata Chalapathi, K. Venkata Ramiah, Proc. Indian Acad. Sci. 1968, 68, 109–122.

[51] A. Dong, X. Ye, J. Chen, Y. Kang, T. Gordon, J. M. Kikkawa, C. B. Murray, J. Am. Chem. Soc. 2011, 133, 998–1006.

[52] A. Nsubuga, M. Sgarzi, K. Zarschler, M. Kubeil, R. Hübner, R. Steudtner, B. Graham, T. Joshi, H. Stephan, Dalton Trans. 2018, 47, 8595–8604.

[53] J. Y. Chane-Ching, A. Lebugle, I. Rousselot, A. Pourpoint, F. Pellé, J. Mater. Chem. 2007, 17, 2904–2913.

[54] K. Liu, X. Liu, Q. Zeng, Y. Zhang, L. Tu, T. Liu, X. Kong, Y. Wang, F. Cao, S. A. G. Lambrechts, M. C. G. Aalders, H. Zhang, ACS Nano 2012, 6, 4054–4062.

[55] F. Ai, Q. Ju, X. Zhang, X. Chen, F. Wang, G. Zhu, Sci. Rep. 2015, 5, 10785.

[56] G. Velazquez, A. Herrera-Gómez, M. O. Martın-Polo, J. Food Eng. 2003, 59, 79–84.

[57] X. Zhao, J. Zhang, L. Shi, M. Xian, C. Dong, S. Shuang, RSC Adv. 2017, 7, 42159–42167.

[58] C. Berthomieu, R. Hienerwadel, Photosynth. Res. 2009, 101, 157–170.

[59] Y. Hu, B. Wu, Q. Jin, X. Wang, Y. Li, Y. Sun, J. Huo, X. Zhao, Talanta 2016, 152, 504–512.

[60] J. Tošović, Kragujevac J. Sci. 2017, 99–108.

[61] Y. Wang, A. Pitto-Barry, A. Habtemariam, I. Romero-Canelon, P. J. Sadler, N. P. E. Barry, Inorg. Chem. Front. 2016, 3, 1058–1064.

[62] M. Baibarac, I. Smaranda, A. Nila, C. Serbschi, Sci. Rep. 2019, 9, 14278.

[63] P. Huang, L. Bao, C. Zhang, J. Lin, T. Luo, D. Yang, M. He, Z. Li, G. Gao, B. Gao, S. Fu, D. Cui, Biomaterials 2011, 32, 9796–9809.

[64] M. Scimeca, S. Bischetti, H. K. Lamsira, R. Bonfiglio, E. Bonanno, Eur. J. Histochem. 2018, 62, 2841.

[65] S. Watanabe, A. Punge, G. Hollopeter, K. I. Willig, R. J. Hobson, M. W. Davis, S. W. Hell, E. M. Jorgensen, Nat. Methods 2011, 8, 80–84.

[66] M. G. Paez-Segala, M. G. Sun, G. Shtengel, S. Viswanathan, M. A. Baird, J. J. Macklin, R. Patel, J. R. Allen, E. S. Howe, G. Piszczek, H. F. Hess, M. W. Davidson, Y. Wang, L. L. Looger, Nat. Methods 2015, 12, 215–218.

[67] M. Kociak, L. F. Zagonel, Ultramicroscopy 2017, 176, 112–131.

[68] F. Wang, R. Deng, X. Liu, Nat. Protoc. 2014, 9, 1634–1644.

[69] C. Siefe, R. D. Mehlenbacher, C. S. Peng, Y. Zhang, S. Fischer, A. Lay, C. A. McLellan, A. P. Alivisatos, S. Chu, J. A. Dionne, J. Am. Chem. Soc. 2019, 141, 16997–17005.

[70] J. F. Tallman, C. C. Smith, R. C. Henneberry, Proc. Natl. Acad. Sci. 1977, 74, 873–877.

[71] L. Xiao, H. Zhu, J. Shu, D. Gong, D. Zheng, J. Gao, Arch. Gynecol. Obstet. 2022, 305, 179–192.

[72] C. Li, A. P. Tardajos, D. Wang, D. Choukroun, K. Van Daele, T. Breugelmans, S. Bals, Ultramicroscopy 2021, 221, 113195.

[73] T. Wagner, J. Eglinger, Thorstenwagner/Ij-Particlesizer: V1.0.9 Snapshot Release, Zenodo, 2017.

