## Supplementary Information for "Cathodoluminescent and Characteristic X-ray-emissive Rare-Earth-doped Core/Shell Immunolabels for Spectromicroscopic Analysis of Cell Surface Receptors"

S. Habermann, Dr. L. R. H. Gerken, Dr. A. Gogos, V. M. Kissling, Prof. Dr. I. K. Herrmann
Laboratory for Particles Biology Interactions, Department Materials Meet Life, Swiss Federal Laboratories for Materials Science and Technology (Empa), Lerchenfeldstrasse 5, 9014 St. Gallen, Switzerland.

Dr. M. Kociak
Université Paris-Saclay, CNRS, Laboratoire de Physique des Solides, Orsay 91405, France.

Dr. C. Monachon
Attolight AG, 1015 Lausanne, Switzerland.

Prof. Dr. I. K. Herrmann*
The Ingenuity Lab, University Hospital Balgrist, Balgrist Campus, Forchstrasse 340, 8008 Zurich, Switzerland.

Prof. Dr. I. K. Herrmann*
Faculty of Medicine, University of Zurich, Rämistrasse 74, 8006 Zurich, Switzerland.

**Keywords:** Nanoparticle, EDX, Ultrastructure, Immunotargeting, Multi-Color

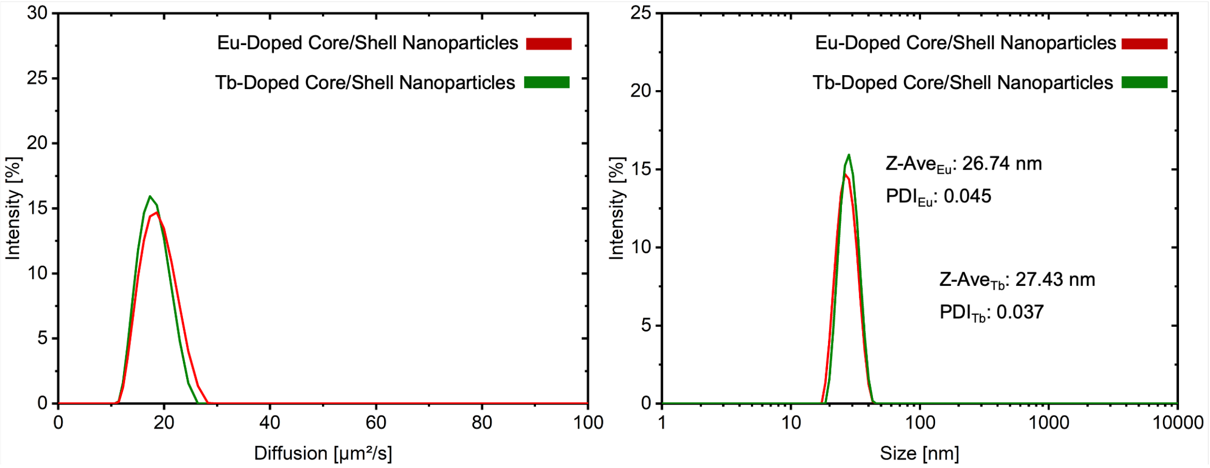


**Suppl. Figure S1:** Diffusional analysis of as-prepared core/shell nanoparticles, revealing an exceptionally narrow size distribution, which foreshadowed nearly identical diffusion properties.

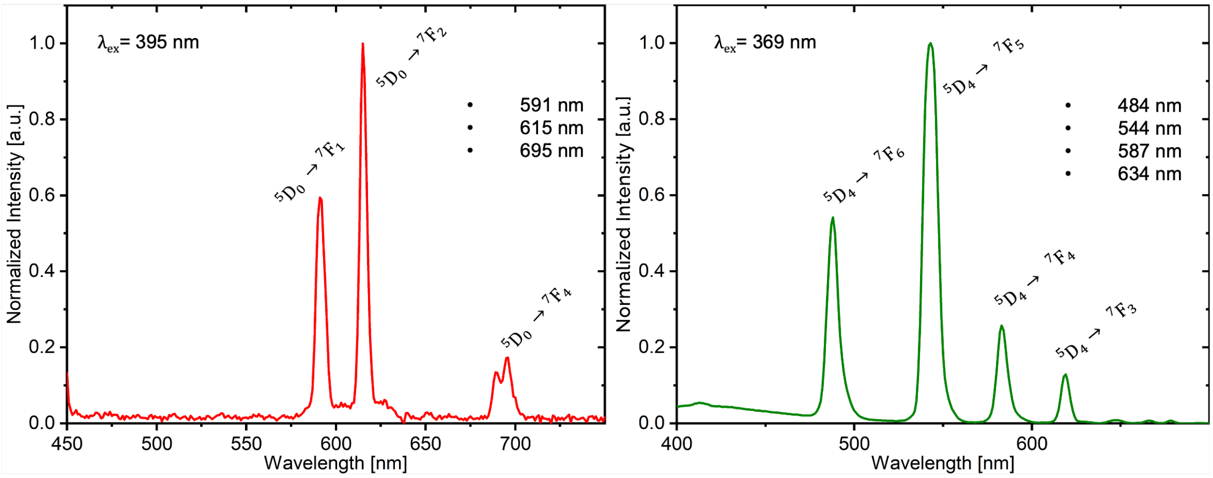


**Suppl. Figure S2:** Photoluminescence spectra of $Eu^{3+}/Tb^{3+}$-doped core/shell $NaGdF_4$ nanoparticles. a.u.: arbitrary units.

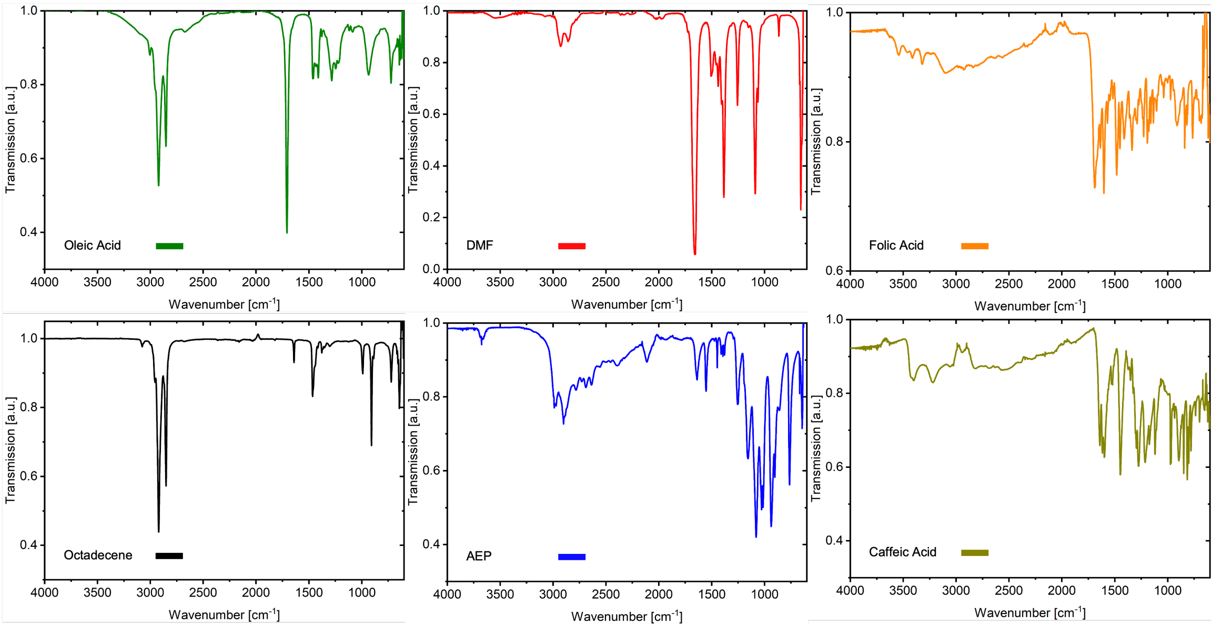


**Suppl. Figure S3:** FTIR analysis of solvents and pure reagents involved during the surface functionalization. a.u.: arbitrary units.

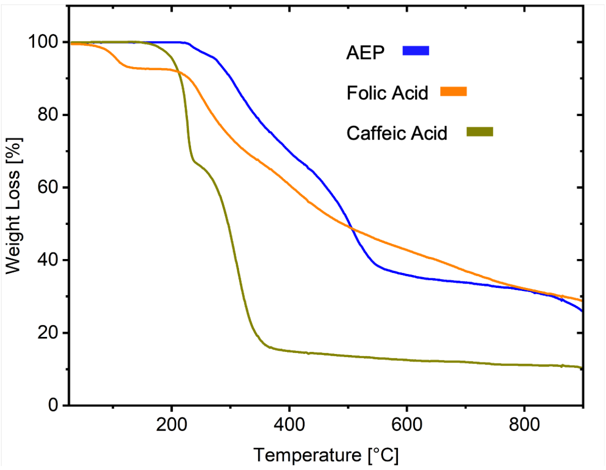


**Suppl. Figure S4:** TGA analysis of the pure surface modifications, used for surface functionalization.

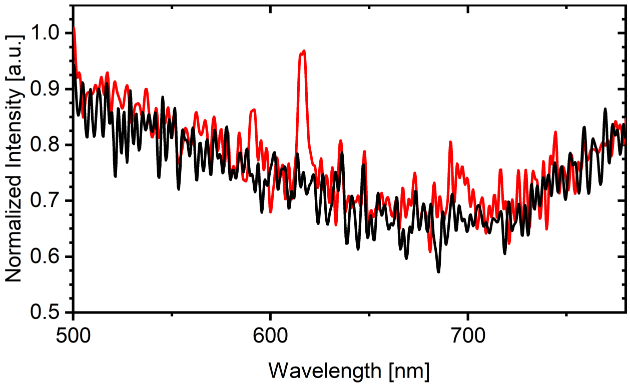


**Suppl. Figure S5:** CL spectrum of europium-doped nanoparticles after simple data processing; 0.366157 Hz Low Pass filter applied. a.u.: arbitrary units.

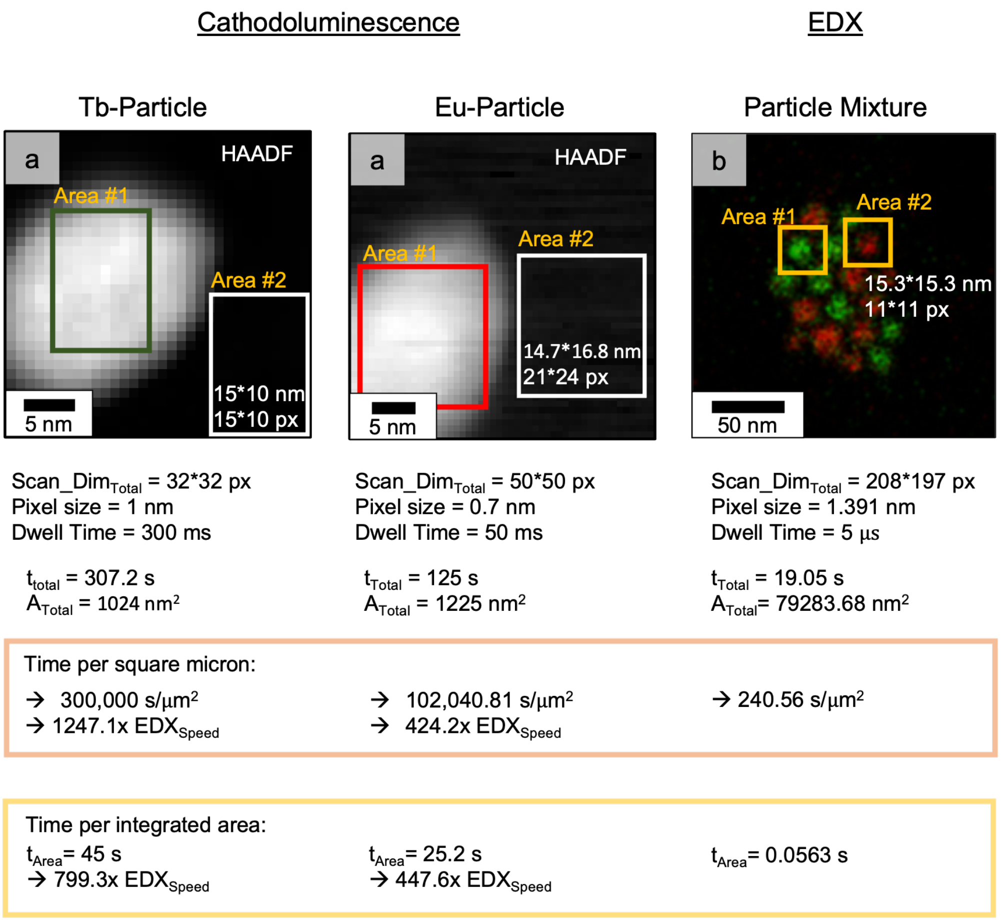


**Suppl. Figure S6:** Comparison of acquisition times used during single-particle analysis. Particularly emphasised are the $t/\mu^2$ (orange) and the time per target (yellow), where the relationships between the experiments are shown in the tables.

The comparison of CL and EDX-SM in Suppl. Figure S6 shows the striking differences in acquisition speed. The CL images were acquired using a higher resolution with a smaller pixel size, however, EDX-SM also offered a resolution suitable for ultrastructural assessment of biological samples. Given a significantly larger area mapped, while requiring a fraction of the time needed for the CL analysis, the potential of EDX-SM is clearly demonstrated. Even if CL is given the benefit of the doubt since the acquisition parameters are not completely consistent, the times per area are of such different magnitudes that EDX-SM showed its advantages, especially when taking into account that within its acquisition time, EDX-SM offered superior data quality. It was therefore assumed that EDX could be assigned a superiority unattainable for CL within this STEM set-up.

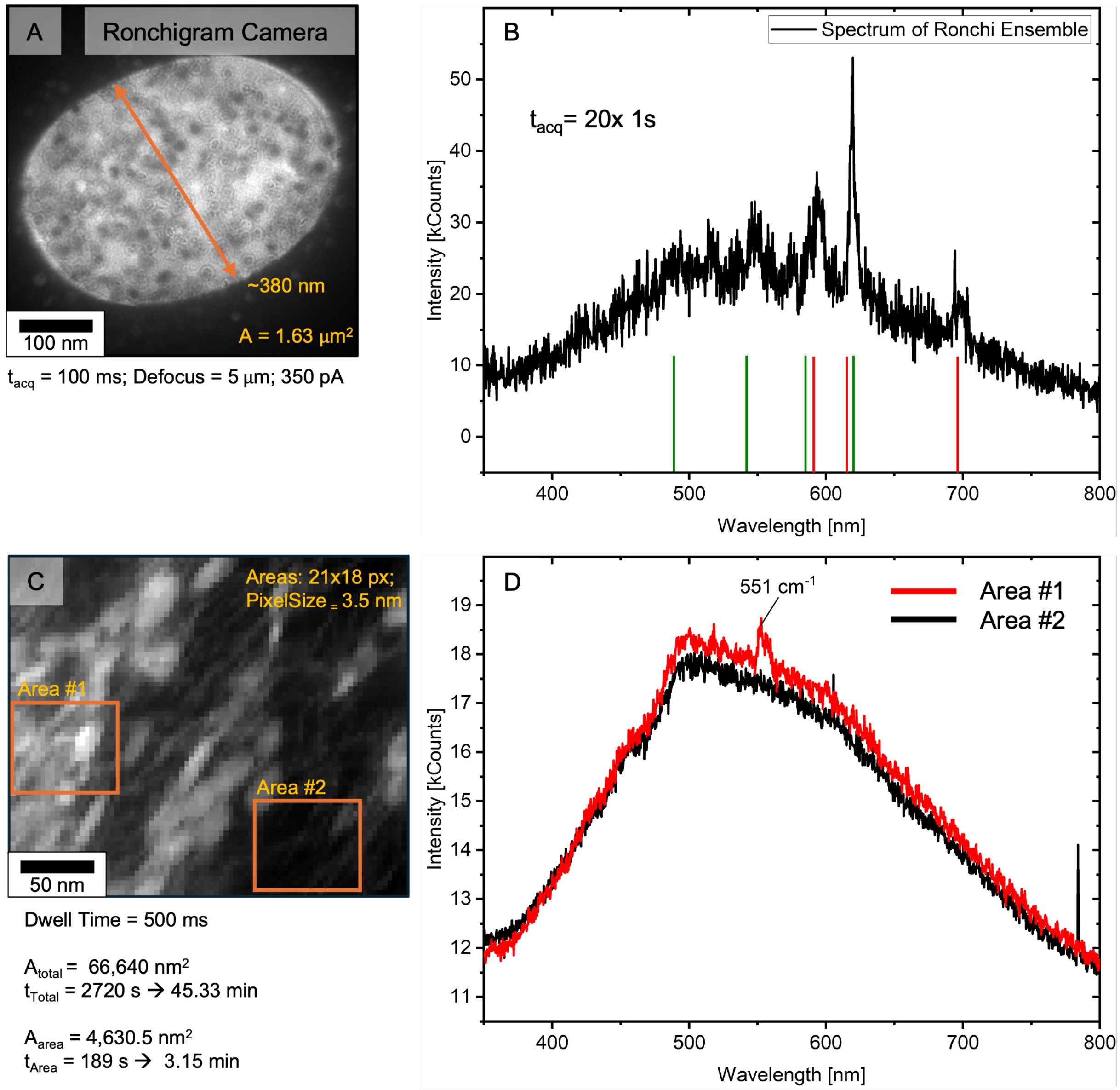


**Suppl. Figure S7:** (**A, B**) Nanoparticle ensemble composed of both particle populations (1:1, weight), selected using the Ronchigram-Cam (**A**) and corresponding CL spectra (**B**) shown. (**C, D**) Tb-doped nanoparticles in cell vicinity. Medium-angle annular dark field (MAADF) given in **C** and CL spectra derived from areas on particles (#1) and on carbon/epoxy (#2) shown in **D**. Areas: 73.5x63 nm (21x18 px). All given particles underwent the particle functionalization and electron microscopy sample preparations as described in section 8.

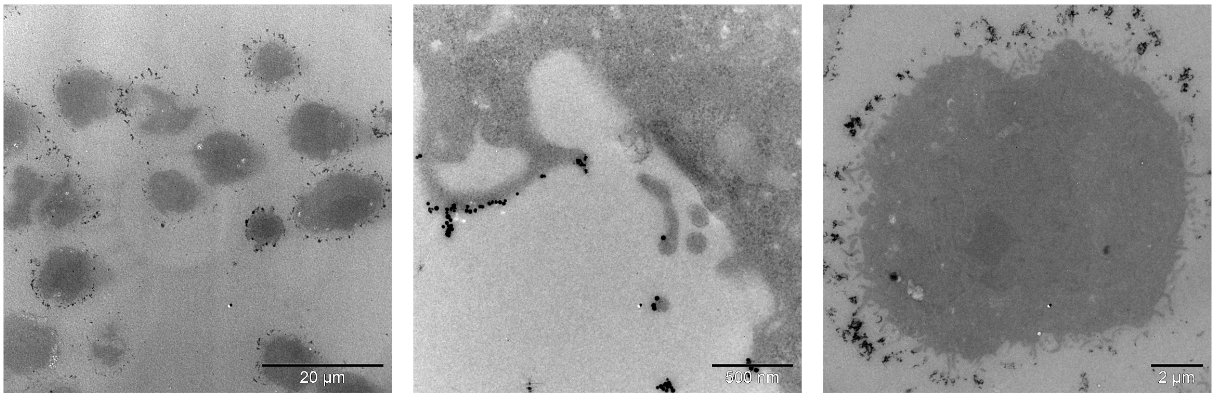


**Suppl. Figure S8:** Bright-field TEM images of 80 nm-sections deposited on formvar grids, showing the binding of the functionalized nanoparticles along the cell membranes without significant unspecific nanoparticle presence in the epoxy or cell vicinity.
